# Accurate *in vivo* population sequencing uncovers drivers of within-host genetic diversity in viruses

**DOI:** 10.1101/349498

**Authors:** Maoz Gelbart, Sheri Harari, Ya’ara Ben-Ari, Talia Kustin, Dana Wolf, Michal Mandelboim, Orna Mor, Pleuni Pennings, Adi Stern

## Abstract

Mutations fuel evolution and facilitate adaptation to novel environments. However, characterizing the spectrum of mutations in a population is obscured by high error rates of next generation sequencing. Here, we present AccuNGS, a novel *in vivo* sequencing approach that detects variants as rare as 1:10,000. Applying it to 46 clinical samples taken from early infections of the human-infecting viruses HIV, RSV and CMV, revealed large differences in within-host genetic diversity among virus populations. Haplotype reconstruction revealed that increased diversity was mostly driven by multiple transmitted/founder viruses in HIV and CMV samples. Conversely, we detected an abundance of defective virus genomes (DVGs) in RSV samples, including hyper-edited genomes, nonsense mutations and single point deletions. Higher proportions of DVGs correlated with increased viral loads, suggesting increased cellular co-infection rates, which enable DVG persistence. AccuNGS establishes a general platform that allows detecting DVGs, and in general, rare variants that drive evolution.

## INTRODUCTION

Viruses are among the fastest evolving entities on earth. Thanks to short generation times, large population sizes and high mutation rates, viruses and in particular RNA viruses rapidly accumulate genetic diversity. This genetic diversity is key to successful adaptation of viruses to novel challenges such as the immune system and drugs (Duffy, et al. 2008). The short time window following virus transmission, termed *acute infection*, is extremely critical for virus populations: they must develop their arsenal of genetic diversity to either escape from the immune system, evade drugs or adapt to a new environment (possibly even a new host) (Parrish, et al. 2008). While we know that many genetic variants are created in this critical time window – the vast majority of these variants segregate at low frequencies that approach the mutation rate of the virus, and are completely undetectable using current next generation sequencing (NGS) approaches (Acevedo, et al. 2014). In fact, these variants will only be discovered after they reach high frequencies, when they will already exert a deleterious effect on the host. For example, today we are only able to detect an HIV drug resistance mutation when it is at a relatively high frequency, after resistance phenotype is observed and already unpreventable (Ram, et al. 2015; Boucher, et al. 2018; Döring, et al. 2018). The naturally occurring frequency of drug resistance mutations could also allow us to infer the fitness cost of these mutations in the absence of drugs (Theys, et al. 2018). An additional long-standing question in the field of virus evolution is the measurement of mutation rates, which also requires accurate sequencing (Acevedo, et al. 2014; Zanini, Puller, et al. 2017). Thus, an accurate sequencing method that maintains high yield is required to thoroughly study evolving populations.

Accurate population sequencing is critical in many different disciplines including genetics, immunology and microbiology as well as tumor screening and prenatal diagnosis. However, using the standard NGS protocols may result in significant background error rates. In fact, following typical post-processing of NGS data, genetic variants that are measured at frequencies lower than 1-5% are discarded (Meacham, et al. 2011; Casadella and Paredes 2017; Huber, et al. 2017; McCrone, et al. 2018). In the past few years, several innovative approaches were suggested to reduce the background error rates of the NGS process: rolling-circle-based redundant coding of the amplified fragments (Lou, et al. 2013; Acevedo, et al. 2014; Reid-Bayliss and Loeb 2017; Wang, et al. 2017); consensus sequencing of barcoded genomic fragments (Jabara, et al. 2011; Kennedy, et al. 2014; Zhou, et al. 2015; Jee, et al. 2016; Newman, et al. 2016; Wang, et al. 2017); error reduction by overlapping paired reads in paired-end sequencing (Chen-Harris, et al. 2013; Schirmer, et al. 2015; Preston, et al. 2016); and usage of improved polymerases (Imashimizu, et al. 2013). However, most experimental methods described above are designed for samples with high biomass and are inapplicable for sequencing of clinical samples, where the biomass may be extremely low. Furthermore, these experimental protocols may introduce their own artifacts to the sequencing process (Lou, et al. 2013; Brodin, et al. 2015). On the computational side, it has been suggested that well-established variant callers do not perform well on clinical virus samples (McCrone and Lauring 2016). Here, we sought to develop a simple and rapid approach that can tackle the problem of accurate sequencing of clinical samples, and applied it to study the early stages of virus infection.

We describe AccuNGS, a simple yet powerful approach for accurate population sequencing and bioinformatics variant calling. We extensively optimize all stages of the method to ensure high accuracy and maximal yield. We use AccuNGS to perform in-depth sequencing of 46 samples from three different major human pathogenic viruses: human immunodeficiency virus (HIV), respiratory syncytial virus (RSV), and cytomegalovirus (CMV), all sampled during the acute infection stage. We compare the within-host genetic diversity among and within different virus populations, and find patterns characteristic of each virus. We demonstrate the role of multiple transmitted/founder viruses as major contributors to the genetic diversity in HIV and CMV. Furthermore, we identify and quantify the impact of various host editing enzymes on the mutational spectrum of viral genomes/populations in vivo. Intriguingly, we find that RSV samples bear much higher levels of potentially defective virus genome (DVGs) than the two other viruses analyzed herein. Finally, we propose a link between the level of DVGs in a population and the levels of cellular co-infection.

## RESULTS

### An overview of the AccuNGS sequencing approach

We have developed a new sequencing approach called AccuNGS, designed to maximize both the accuracy of variant calling and template recovery. To achieve this, several key concepts were put together: (i) usage of high-yield and high-fidelity polymerases to increase yield and reduce error rates; (ii) sequencing error reduction through overlapping paired end reads; (iii) minimization of template loss across different stages of the protocol, (iv) statistical modeling of error rates using a control that forms the heart of a computational pipeline for variant calling and inference of variant frequency (Fig. 1). We first extensively optimized the different experimental stages of the protocol, by testing the accuracy of sequencing a clonally derived plasmid (Methods). This allowed us to assess the contribution of the various stages to the error rate of the protocol (Supplementary text, Fig. S1, Tables S1-S5). We were able to (a) mostly rule out the contribution of PCR to error rates (Zanini, Brodin, et al. 2017), (b) note that the two reads generated by the sequencer are dependent observations, and (c) reveal that AccuNGS is likely limited by the accuracy of the sequencing machine itself (which we cannot affect) (Supplementary text, Fig. S1B). We conclude that the mean error rate for transitions and transversions that is achieved by AccuNGS is a little lower than 10^−4^ and 10^−5^, respectively, and varies by types of mutation (Table 1). This allows us to statistically call variants that are as rare as 10^−4^, which we further verified by creating synthetic mixtures of two different HIV plasmids at different proportions and sequencing them (Fig. S2).

**Table 1.**
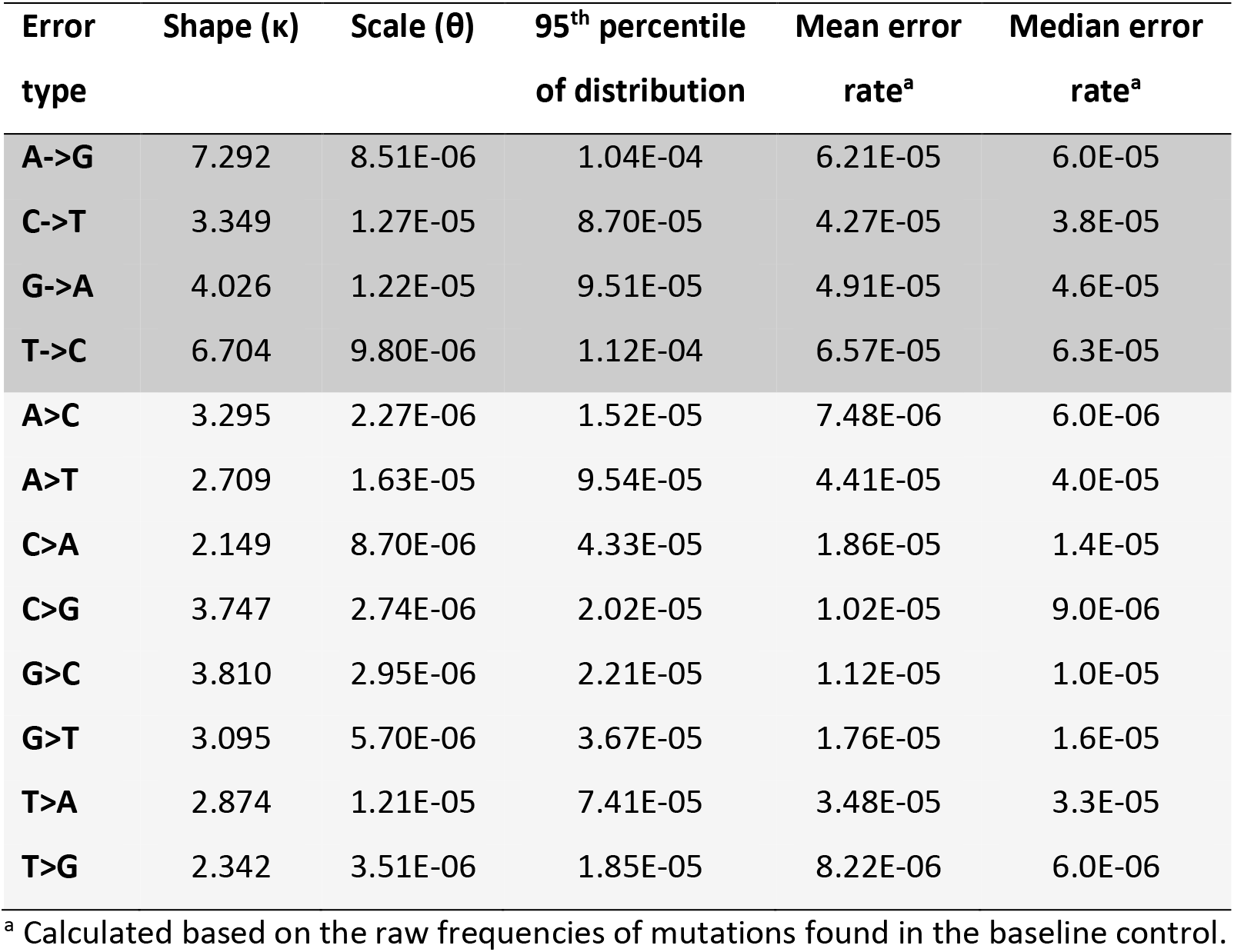
Error rates for AccuNGS. Shown are parameters for fitted gamma distributions given a Q30 score cutoff. Transitions are shaded in darker gray than transversions.

**Fig. 1.**
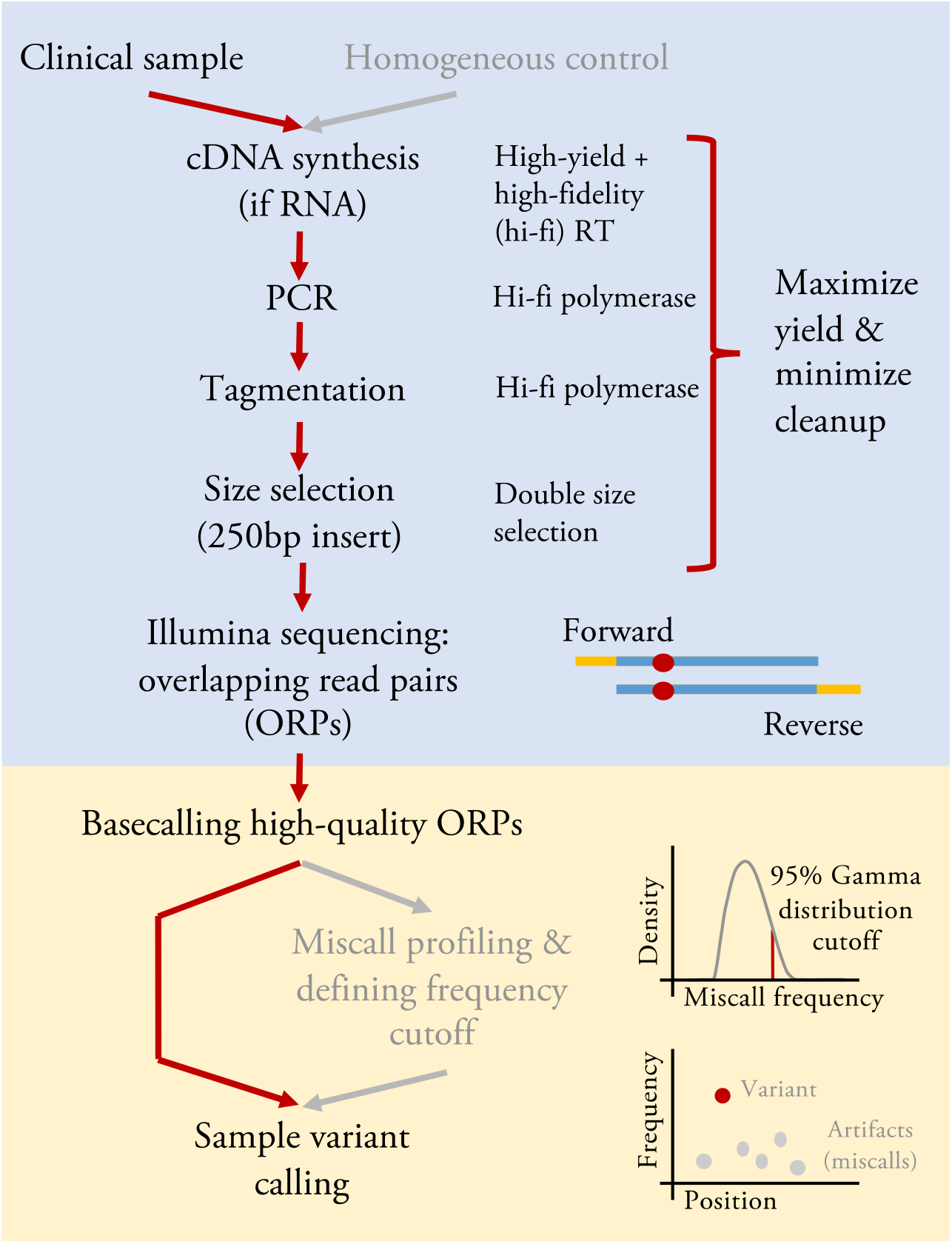
AccuNGS principles. The protocol requires side by side targeted sequencing of a biological sample together with a homogenous control (e.g., a plasmid encoding for the same genomic region sequenced in the sample). Stages of the protocol include: (i) High-fidelity and high-yield RT reaction, (ii) High-fidelity PCR reaction, (iii) Tagmentation and library construction with size selection for an insert the size of a single paired-end read, (iv) Paired-end sequencing where each base in the insert is sequenced twice, once in the forward read and again in the reverse read, (v) Alignment of reads, Q-score filtering on both reads and base-calling of both the sample and a homogeneous control, and (vi) Sample variant calling based on gamma distributions that are fitted to the process errors, defined as mutations found in the homogeneous control.

Initially we used a barcoding approach (Jabara, et al. 2011) to estimate the number of viral genomes we sequenced for one of the HIV clinical samples, allowing us to estimate that we had sequenced ~15,000 separate genomes (Supplementary text). While it was reassuring that we sequenced a large number of genomes, we also showed that the mere addition of a barcode led to a reduction in the number of sequenced templates (Supplementary text, Fig. S3). In other words, we found that without a barcode we sequence more genomes, but we cannot count how many. We henceforth avoided the use of barcoding. Finally, we would like to emphasize that while the *average* error rate of AccuNGS is very low, sampling and PCR biases may lead to a given variant being detected at a lower or higher frequency than it should be (Illingworth, et al. 2017; Zhao and Illingworth 2019). This is especially true when the number of input templates is low. We thus conclude that AccuNGS is useful for inferring average rates and average frequencies (e.g., premature stop codons as elaborated below) rather than inferring the exact frequency of one given variant in a given sample.

### Accurate sequencing of different virus populations during acute/early infections

We obtained a total of 46 samples from patients recently infected by the RNA viruses HIV and RSV, and the DNA virus CMV (Table 2, Table S6). We focused our sequencing efforts mostly on conserved genes, precisely since we expect less diversity and we wanted to test our ability to detect rare “hidden” variants that are otherwise unobservable. Hyper-variable genes such as the HIV-1 envelope have been sequenced extensively using other sequencing approaches (e.g., Keele, et al. 2008; Salazar-Gonzalez, et al. 2008; Zhou, et al. 2016), and the presence of high frequency variants is not surprising in such genes. We thus chose the Gag-Pol open reading frame for HIV, the M2 and L open reading frame (encoding for the viral polymerase) for RSV, and the UL54 (also encoding for the viral polymerase) for CMV. In order to allow for comparison, we also sequenced the F and G envelope glycoproteins genes in RSV, and further compared our results to previous sequencing results of the envelope gene in HIV.

**Table 2.** Details of samples sequenced from clinical virus samples.

| Virus | #Samples | Samples' origin | Estimated weeks post infection | Region sequenced (NCBI ID) |
| --- | --- | --- | --- | --- |
| HIV | 9 | Plasma | 2-5 | 532:3280 (K03455) |
| RSV | 25 | Nasal/throat swabs | <1 | 4640-14350 (U39661) |
| CMV | 12 | Amniotic fluid / Urine / saliva | >=4 | 78200-81912 (NC_006273) |

Each sample underwent population sequencing and variant calling using AccuNGS (see Methods). We began by calculating the total nucleotide diversity π in each sample based on the transition variants (Methods). This revealed different distributions of diversity within and between viruses (Fig. 2A). In the HIV populations, diversity values spanned several orders of magnitude. On the contrary, RSV populations exhibited very similar intermediate levels of diversity. Similarly, CMV populations usually displayed the lowest diversity with the exception of one sample. We set out to understand what factors drive the differences in diversity among the different samples.

**Fig. 2.**
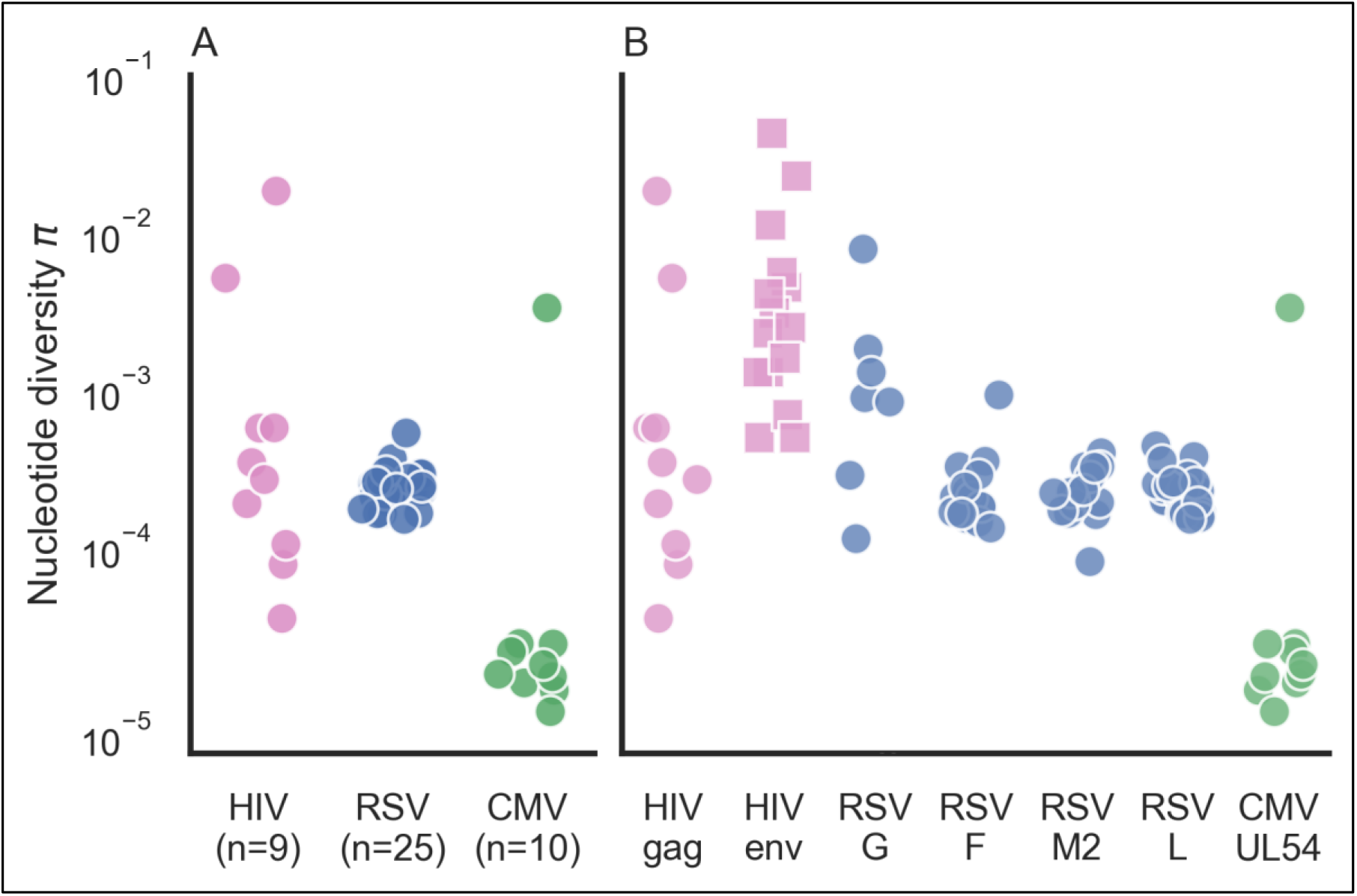
Nucleotide diversity π for acute infections across different virus samples and genes. (A) Each point represents the π diversity of a single sample, across all genes sequenced. Diversity values were calculated using transition mutations only. (B) Gene by gene breakdown of nucleotide π diversity. Values for HIV envelope (squares) were taken from previously published data (Salazar-Gonzalez, et al. 2008).

### Mutation and selection

We first considered the two most evident evolutionary causes of differences in diversity: mutation and selection. First, when considering the mutation rate of a virus, it is clear that the only DNA virus in our data has much lower mutation rates than RNA viruses (Sanjuan, et al. 2010) and indeed displays lower diversity. The two RNA viruses display more diversity than the DNA virus CMV, but the variation in diversity levels is much higher in HIV than in RSV.

We further considered whether differences in selection pressure cause the variation in diversity we see among the RNA virus samples. This was unlikely to cause within-virus differences, since we sequenced the same set of genes within each of the virus samples. We did note that the immunogenic envelope proteins in this study (HIV Env and RSV G proteins), known to be under positive selection (Seibert, et al. 1995; Nielsen and Yang 1998; Tan, et al. 2013), displayed on average higher diversity than the conserved genes (Fig. 2B). This is not surprising given that some mutations in envelope proteins will be under positive selection since they allow immune evasion, and hence may reach higher frequencies. However, this could not explain why we saw dramatic differences in diversity in different samples from the same virus when focusing on the same gene (e.g., *gag* in HIV).

### Transmission bottleneck size as a contributor to genetic diversity during acute infections

It has previously been noted that infections initiated by a few different divergent viruses are characterized by higher genetic diversity (Keele, et al. 2008; Cudini, et al. 2019). Visual inspection of our frequency plots (Fig. 3, Fig. S4-S6) suggested that often variant frequencies were strongly imbalanced, also evident as “bands” of variants at similar frequencies. For example, sample HIV6 (measured diversity 1.46×10^−2^) contained many variants segregating at a frequency of ~2×10^−1^ yet very few variants segregated at frequencies between 10^−3^ and 10^−1^ (Fig. 3A). We first considered how likely it is that such a sample would be initiated by only one founder virus/genotype, where all variants begin at a defined frequency of zero. Given a large enough population size and a mutation rate in the order of 10^−5^ mutations/site/day (Zanini, Puller, et al. 2017), we expect neutral variants that are likely generated over and over almost every day to roughly reach a frequency of 10^−4^-10^−3^ after a few weeks of infection, which is much lower than 10^−1^. Genetic drift or positive selection could drive a few variants to increase in frequency over a short time; however, it seems extremely unlikely that there is such a large set of sites under the exact same regime of positive selection, especially as we had sequenced a gene where positive selection is all in all less prevalent, at least this early in the course of the infection. Thus, it seems quite unlikely that very high diversity samples containing many high frequency variants are founded by one virus genotype, and a more likely explanation is the presence of multiple transmitted/founder viruses.

**Figure 3.**
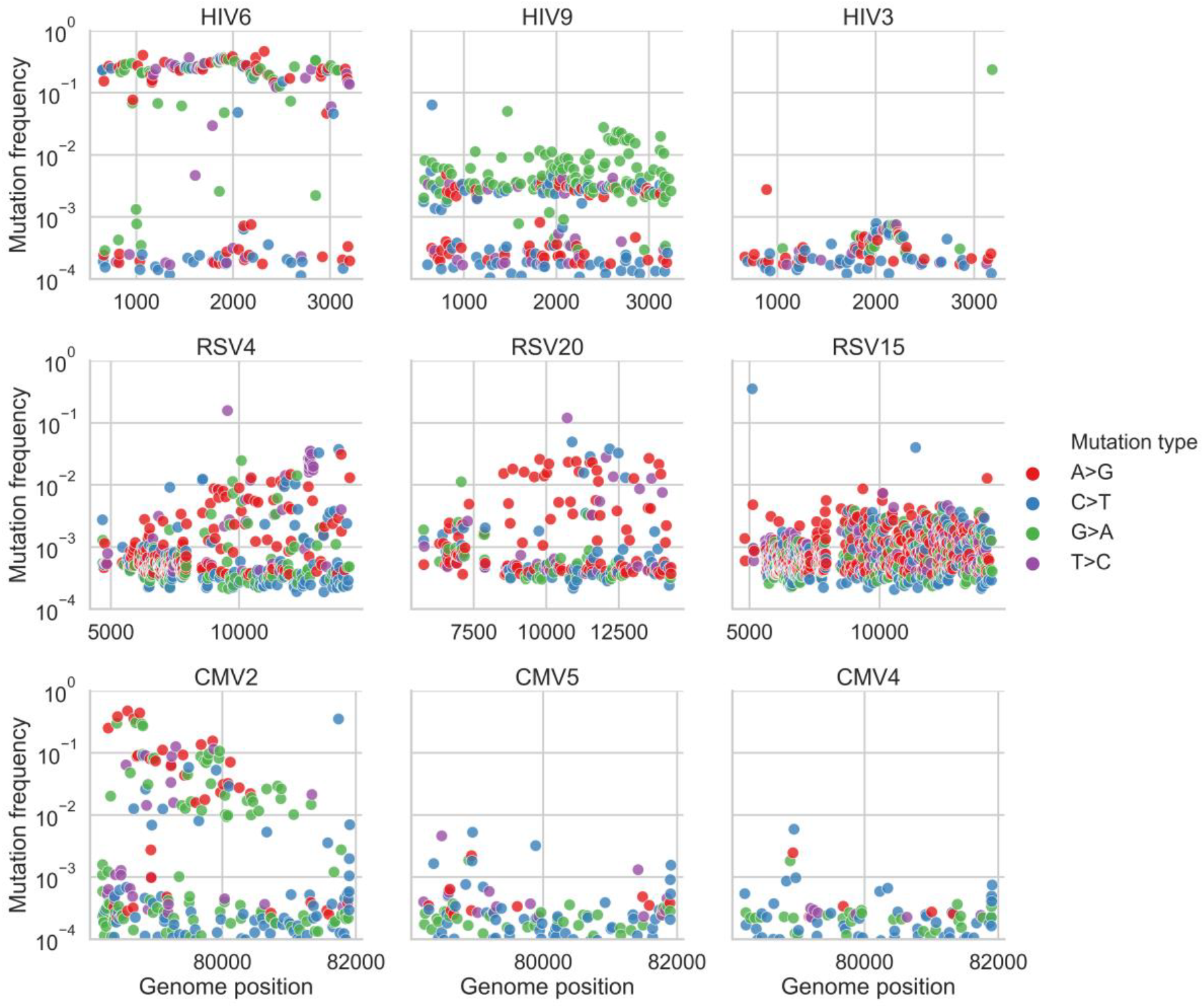
Variants frequency plots in representative samples. Shown are frequencies of transition variants called by AccuNGS, for representative samples from each virus (HIV, top row, RSV, middle row, CMV, bottom row). Samples exemplify multiple founder infections, mutation biases, and relatively homogenous populations (see text for details).

### Inferring haplotypes and multiple founders

To evaluate the number of founder viruses we require an estimation of the different haplotypes present in a sample, and their abundances. However, reconstruction of virus haplotypes from short reads and from one time-point is a longstanding problem (Schirmer, et al. 2014). This is due to two conflicting features of viral population sequencing data: on the one hand, the data is often too homogenous. In other words, most reads are identical or almost identical to the consensus, and there may not be enough variants on one read that allow “linking” it with another read. On the other hand, the mutation rates of viruses may scale with sequencing error rates, throwing off most commonly used haplotype reconstruction methods.

Two major advantages of our data are the much higher accuracy of AccuNGS, coupled with very high sequencing depth. We thus set out to develop a new approach for inferring viral haplotypes. Instead of attempting to reconstruct the entire haplotype, we mainly focused on inferring if more than one haplotype (and hence more than one founder virus in our case) is present in a sample. Our approach is based on looking for statistical enrichment for two variants being present on the same read as opposed to each variant on its own, and then linking these reads one with another based on shared variants (Methods) (Yang, et al. 2013). To validate our approach, we used the samples where we had synthetically mixed two plasmids and discovered that all and only true variant sites were identified as linked to each other in with an accurately estimated frequency of 1:10,000 (Fig. S7). Importantly, this confirmed that AccuNGS does not suffer from various artifacts such as PCR recombination that could break linkage between adjacent sites, and that even a rare haplotype can be inferred.

Our haplotype reconstruction approach also led us to realize one of the combined strength and pitfalls of accurate sequencing: we were able to initially detect minute contaminations (a few hundred out of millions of reads) from one sample into another, which we were then able to computationally filter out (Supplementary text). This contamination likely occurred during one of the stages of the library preparation and sequencing, and emphasizes the sensitivity of genomics approaches today, which may often exceed that of the molecular biology itself (Kircher, et al. 2011; Gu, et al. 2019). On the other hand, AccuNGS also allows for the clear-cut detection and evaluation of any contamination, which we believe are very important to capture.

We next applied our haplotype inference flow to all the filtered samples, and found that all of the high diversity samples (total diversity >10^−3^, Fig. 2A) exhibited strong evidence for containing two or more divergent haplotypes (Figs. S4-S6). Two examples are shown in Figs. 3 and 4: HIV sample 6 has a “band” of variant frequencies around 2×10^−1^ (Fig. 3), and indeed most of these variants can be linked to each other in this sample (Fig. 4). CMV sample 2 has a wide “band” of variant frequencies between 10^−2^ and 5×10^−1^ (Fig. 3), which were also mostly found to be linked, and likely represent a founder haplotype and the associated variants that were created on the background of this haplotype (Fig. 4). In general, we found no evidence for two or more haplotypes in the less diverse samples, except for the most diverse RSV sample that also showed limited evidence of a low frequency haplotype (Fig. S5) (see discussion).

**Figure 4.**
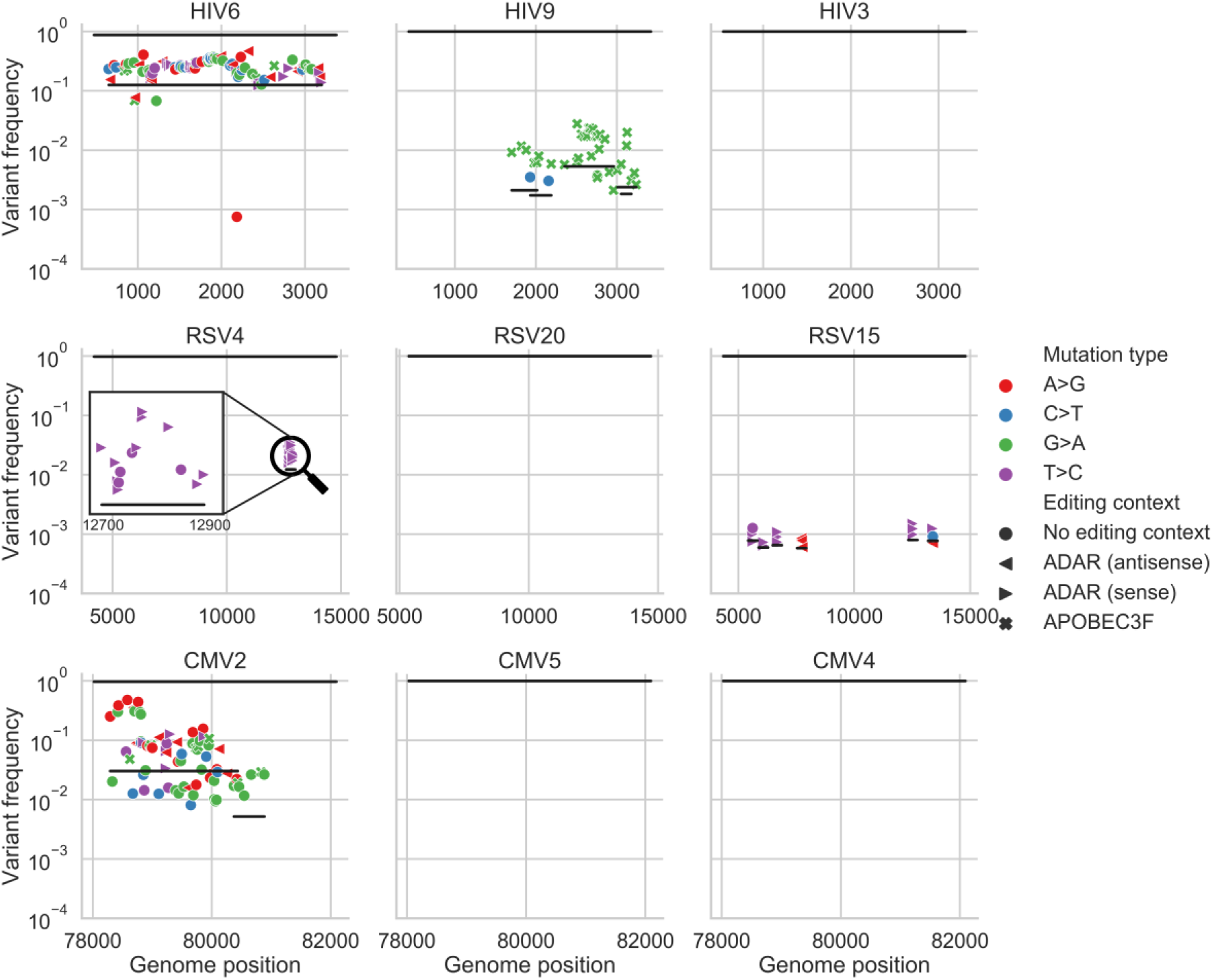
Haplotype reconstruction based on co-occurrence of variants on the same reads. Shown are inferred haplotypes (lines) based on consecutive significant associations of pairs of variants (shapes) one to another on the same read. The uppermost line in each panel represents the consensus sequence, which by definition is the major haplotype in each sample. Both HIV6 and CMV2 samples show strong evidence of an additional haplotype, which is likely a second founder genotype. Sample HIV9 shows evidence of G>A hyper-mutation in the context of APOBEC3 editing, samples RSV4 and RSV15 show evidence of T>C or A>G hyper-mutation in the context of ADAR editing in regions spanning a few hundred bases. The hyper-mutated region in RSV4 sample is magnified for clarity. “Empty” panels signify what are likely single haplotype infections, with no evidence of hyper-mutation.

### Short hyper-mutated genomic stretches

One well-known phenomenon of HIV infections is the potential of host APOBEC3 (A3) proteins to induce hyper-editing on the negative strand of nascent HIV DNA during reverse transcription, resulting in an excess of G>A mutations in regions of the RNA genome (Hache, et al. 2006; Malim 2009). This hyper-mutation strategy is thought to lead to DVGs that are unable to replicate. However, HIV encodes for a gene called *vif* that counteracts A3 proteins, and thus most HIV viruses sequenced from blood samples show only minor evidence for A3 activity (Cuevas, et al. 2015). Similarly, the family of human ADAR proteins have also been shown to induce A>I mutations (read as A>G mutations) in a variety of viruses (Samuel 2012). We set out to test if we detect signals of hyper-editing in our samples. In particular we sought to find stretches of hyper-mutations using our haplotype reconstruction approach in order to evaluate whether hyper-editing contributes to the observed genetic diversity, and to what extent.

Of all 46 samples, only one HIV sample (HIV9, Fig. 3) displayed strong evidence for G>A hyper-editing. In this sample, editing seemed to be widespread, with multiple distinct and overlapping hyper-mutated genomes (Fig. 4). Hyper G>A mutations were enriched in the context of GpA which is the APOBEC3D/F/H favored editing context but not of the canonical APOBEC3G (Beale, et al. 2004; Bishop, et al. 2004). Most variants on these hyper-mutated stretches were missense variants; some of these stretches contained variants that lead to premature stop codons which are presumably lethal for the virus (Fig. S4). The maximum frequency of such variants in the sample was roughly 2×10^−2^. To test whether this occurs due to an inactive *vif* gene, we sequenced this gene in this sample using AccuNGS. We found no support for this hypothesis since the consensus sequence of this gene was intact, but we once again noticed a relatively high level of G>A mutations in the *vif* gene itself (Fig. S8). Notably, this sample was taken from the patient with the highest viral load (VL) in our study, which was >10 million cp/ml.

Out of 25 independent RSV populations, 11 (44%) exhibited evidence of ADAR-mediated hyper-edited genomes, manifested as at least three ADAR-associated mutations on the same haplotype (Whitmer, et al. 2018). The presence of ADAR-like hyper-edited haplotypes was positively correlated with viral loads (R^2^=0.59; logistic regression; p<0.0001), for example all 9 samples with measured VL>650k cp/ml contained hyper-mutated inferred haplotypes (Fig. 5D). When observed, ADAR-like linked variants were present at frequencies varying between ~10^−3^ and ~10^−2^, which by far exceed the mutation rate of any known virus. Out of 26 ADAR-like hyper-edited haplotypes, 23 of them were on the negative strand and only 3 were on the positive strand, in line with previous studies demonstrating that most ADAR-like mutations are acquired on the negative strand of (-)ssRNA viruses (e.g., Whitmer, et al. 2018). Most of the ADAR-like variants on these stretches were missense variants, suggesting they have a detrimental effect on the virus (Fig. S5).

**Figure 5.**
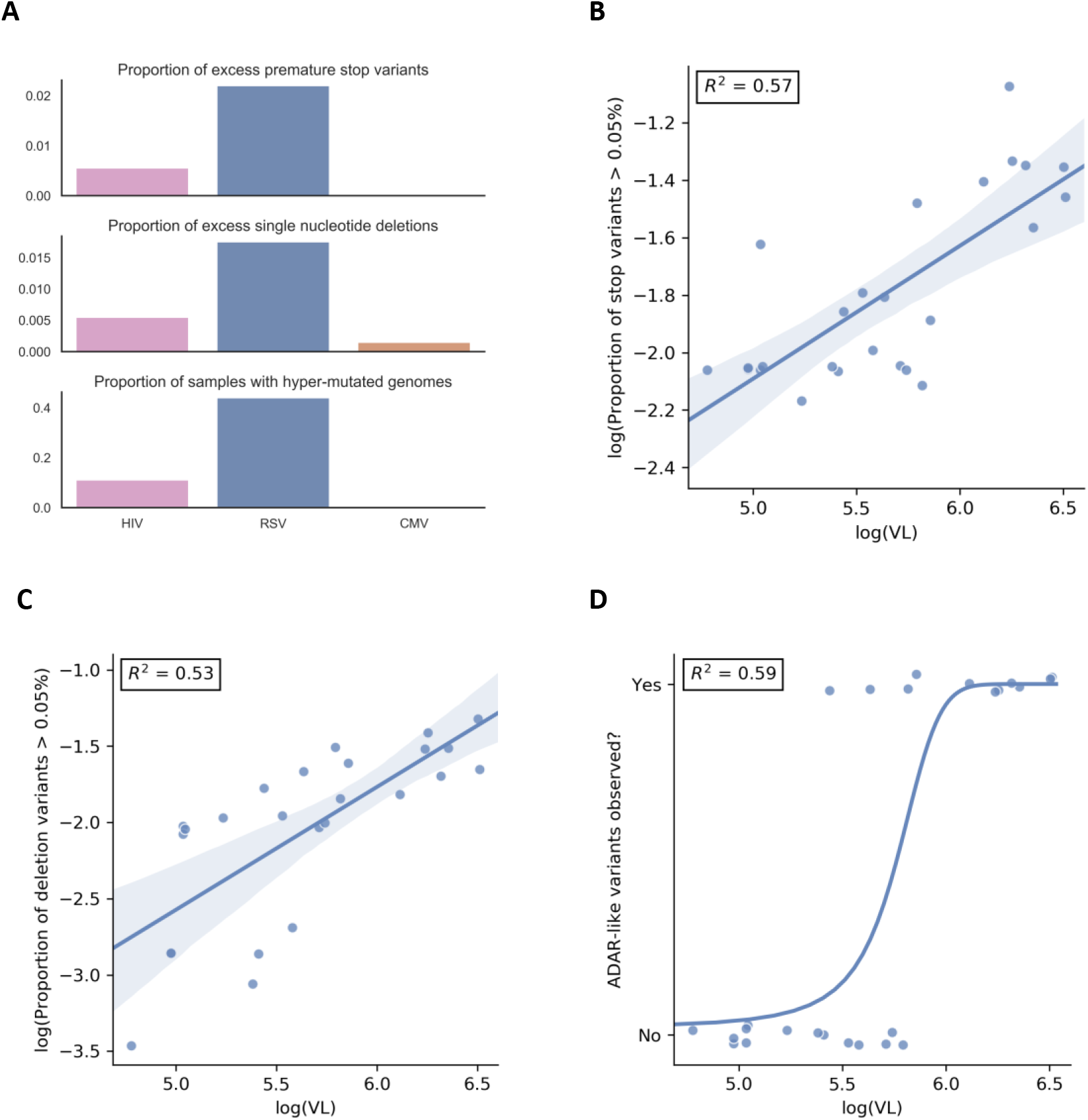
RSV contains high levels of DVGs. (A) Proportion of sites across all samples identified as containing excess lethal variants (top and middle panels), proportion of samples containing at least one hyper-mutated haplotype with a frequency above 5×10^−4^ (bottom panel). For RSV, viral load (VL) positively correlated with the (B) excess premature stop codons, (C) excess single nucleotide deletions, and (D) presence of ADAR-like hyper-edited stretches. An excessive lethal variant is defined as one exceeding a frequency of 5×10^−4^. (B) and (C) display linear regression lines, whereas (D) displays a logistic regression line.

None of the CMV populations exhibited any A3, ADAR, or other pattern of hyper-mutation, suggesting that these hyper-mutating enzymes do not act on CMV populations, at least not for the gene sequenced here, or at the level of detection of AccuNGS (but see Weisblum, et al. 2017).

### Defective virus genomes

Under mutation-selection balance, lethal variants, which are the ultimate DVGs, are expected to segregate at the frequency at which they are generated, which is the mutation rate (Acevedo, et al. 2014; Cuevas, et al. 2015). Based on the expected mutation rates in the viruses we sequenced (<=10^−4^ for HIV and RSV, and ~10^−7^ for CMV (Sanjuan, et al. 2010)), our initial expectation was that lethal variants should rarely be observed. We were therefore surprised to note the existence of premature stop codons, one of the most obvious forms of a lethal mutation, at a frequency of 10^−3^ - 10^−2^ in some of the RSV samples (Fig. S5, Fig. S8). All in all we found that RSV substantially differed from the two other viruses with respect to three markers of excess lethal variants: high frequency premature stop codons, high frequency single nucleotide deletions, and presence of hyper-edited haplotypes. RSV had higher proportions of all three types of lethal mutations (Fig. 5A). It thus appears that mutation-selection balance does not necessarily hold for RSV (at least, for some RSV infections). Similar to the presence of hyper-edited haplotypes for RSV, an excess of stop variants and deletion variants in RSV positively correlated with viral load (VL) (Fig. 5B-D). This raises the possibility that extensive complementation occurs in some RSV samples and in the HIV sample described above, allowing the “rescue” of DVGs through high rates of co-infection.

## DISCUSSION

Application of next generation sequencing to clinical samples is still limited by the ability to reliably capture minor variants (Clutter, et al. 2016; McCrone and Lauring 2016; Boucher, et al. 2018). Here we describe AccuNGS, a simple, rapid and accurate experimental protocol and associated computational pipeline for detecting ultra-rare variants from low-biomass clinical RNA and DNA samples. AccuNGS aims to accurately detect minor variants present in a population of genomes at frequencies of 1:10,000 or lower, close to the mutation rate of RNA viruses (Sanjuan, et al. 2010).

We used AccuNGS to characterize HIV, RSV, and CMV diversity. The HIV-1 samples were from early stages of infection, typically 2-5 weeks post infection, based on serology testing. The RSV samples were taken from children hospitalized due to respiratory problems, about 3-5 days post infection (Lessler, et al. 2009), while the CMV samples taken from amniotic fluid or newborn urine and saliva are several weeks post infection. Most previous studies have reported a nearly completely homogenous virus population during early stages of infection (Henn, et al. 2012; Zanini, et al. 2015; Kijak, et al. 2017). Nevertheless, in all samples sequenced, AccuNGS captured dozens to thousands of minor transition and transversion variants, mostly segregating at frequencies between 1:100 and 1:10,000 (Figs. S4-S6). These variants are most likely the genetic reservoir that serves to fuel the future evolution of these virus populations.

Our results suggest that a prominent factor in determining the intra-host genetic diversity of a sample during acute infection is the number of founder viruses, and in fact we find that the samples with the highest levels of diversity always show evidence for multiple founder infection. A debate has arisen regarding the diversity in CMV samples, where one study has claimed that diversity in this DNA virus is comparable to diversity in RNA viruses (Renzette, et al. 2011), and others suggest that diversity is low in single founder infections and is elevated only when multiple founders/genotypes initiate the infection (Cudini, et al. 2019). Our results strongly support the latter hypothesis, and in fact the high resolution of AccuNGS suggests even lower diversity for single founder new CMV infections than what has been previously reported (Cudini, et al. 2019).

For RSV samples, we found no evidence for multiple haplotypes in the samples, which is somewhat surprising given that this is an airborne virus that reaches high titers. Previous work has suggested that the number of founders in RSV infections is 25±35 (Lau, et al. 2017). Notably, these values were obtained for adults experimentally inoculated with RSV, whereas our study represent natural infection of infants. However, another explanation for this discrepancy is that our data does not allow us to detect infection with multiple founders when they share very similar genotypes, since our haplotype reconstruction method relies on detecting short reads that share two or more mutations. Given the very short duration of RSV infection, it is possible that relatively little genetic diversity is created do novo, and hence very little genetic diversity is transmitted. In other words, an infection may be initiated by several very genetically similar founder genotypes, but we would not detect it. On the other hand, CMV and HIV create longer infections, and the potential to generate and transmit more diverse genotypes within a single carrier is higher.

Our results enabled pinpointing the activity of viral hyper-editing by host enzymes, namely APOBEC3 enzymes and ADAR. The latter was particularly prominent in the RSV infections, where we found distinct clusters of mutations matching ADAR context. Surprisingly, the frequency of these clustered mutations was often relatively quite high, as discussed above. There is a debate today surrounding the role of ADAR in viral infections: in some case it was found to be pro-viral whereas in other cases it has been shown to be anti-viral. Pro-viral activity may be plausible when considering that it has been found that ADAR protects cellular transcripts from being detected by intracellular innate immune response (Liddicoat, et al. 2015; Pfaller, et al. 2018).

Is it possible that the ADAR signatures we find represent edited viral genomes that escape innate immunity? If so, this would mean these are not DVGs but rather haplotypes with a selective advantage. We consider that this is unlikely: many of the ADAR-like mutations we find are non-synonymous, with often 5-10 such mutations found in a short region. It is highly improbable that so many mutations would yield a “viable” genome, and we hence conclude that ADAR-like hyper-editing yields DVGs. This comes together with evidence for other DVGs (premature stop mutants and point deletions) that also segregate at similar frequencies in the high viral load RSV samples. We find that the most likely explanation for this phenomenon is that the rate of cellular co-infection is very high in some RSV samples, which may be promoted by the syncytia that RSV creates, allowing for complementation of these DVGs. In fact, given rough estimates of RSV mutation rates around 10^−4^-10^−5^, we are able to infer that the rate of co-infection should be between 0.9 and 0.99 in some of our RSV samples (Wilke and Novella 2003), which is clearly very high. Moreover, we observe that samples with more DVGs are samples with higher viral loads. Putting this together, we suggest that RSV infections probably occur in a relatively dense site, which allows for so many co-infections, and for the propagation of DVGs.

The relationship between viral loads and the levels of DVGs in a sample is quite intuitive. This suggests that in many cases, DVGs may be incorrectly neglected in downstream analyses. For example, the assumption that such mutations occur at mutation selection balance is used to estimate mutation rates and/or fitness costs (Acevedo, et al. 2014; Cuevas, et al. 2015; Theys, et al. 2018), yet high levels of co-infection may lead to deviations from this balance. We suggest that the use of AccuNGS can detect the rate at which DVGs occur, allowing the inference of the co-infection rate. More generally, AccuNGS has the power to uncover the presence and rate of rare genomic events, such as mutations and editing, directly from clinical samples, allowing a better and more detailed understanding of the processes that govern evolution.

## MATERIALS AND METHODS

### Ethics declaration

The study was approved by the local institutional review boards of Tel-Aviv University, Sheba Medical Center (approval number SMC 4631-17 for HIV and SMC 5653-18 for RSV), and Haddasah Medical Center (approval number HMO-063911 for CMV).

### Reagents and Kits

Unless stated otherwise, all the described reactions in this paper were carrier with the described products according to the manufacturer’s instructions: gel purifications were performed using Wizard® SV Gel and PCR Clean-Up System (Promega, Madison, WI, USA); beads purifications were performed using AMPure XP beads (Beckman Coulter, Brea, CA, USA); concentrations were determined using Qubit fluorometer (Thermo Fisher Scientific, Waltham, MA, USA); reverse transcription (RT) reactions were performed using SuperScript III or IV Reverse Transcriptase (Thermo Fisher Scientific); polymerase chain reactions (PCR) were made using Platinum™ SuperFi™ high-fidelity DNA Polymerase (Thermo Fisher Scientific) or Q5 high-fidelity DNA Polymerase (New England Biolabs (NEB), Ipswich, MA, USA).

### Generation of amplicons from HIV-1 clinical samples

#### Clinical HIV-1 samples

Plasma samples from nine recently diagnosed HIV-1 patients with viral loads of 5×10^5^-1×10^7^ cp/ml were provided by the National HIV Reference Laboratory, Chaim Sheba Medical Center, Ramat-Gan, Israel (Table S6). HIV-1 viral loads were determined and RNA extracted from 0.5 mL as described above. From each sample a maximum of 300,000 HIV-1 copies were reverse transcribed using random hexamer priming.

#### HIV-1 control sample

A pLAI.2 plasmid was linearized using the SalI restriction enzyme (NEB) according to manufacturer’s instruction. In order to create RNA a PCR reaction was set up using primer containing the sequence of the T7 RNA polymerase promotor 5’TAA TAC GAC TCA CTA TAG CTG GGA GCT CTC TGG CTA AC and the primer pLAI 5761-5782 5’GAG ACT CCC TGA CCC AGA TGC C using Q5 DNA polymerase and the following PCR program: initial denaturation for 3min at 98°C, followed by 40 cycles of denaturation for 10sec at 98°C, annealing for 30sec at 65°C and extension for 3min at 72°C, and final extension for 5min at 72°C. Eight microliters from PCR reaction were carried to in-vitro transcription reaction using HiScribe™ T7 High Yield RNA Synthesis Kit (NEB) according to the manufacturer’s instruction. Finally, the RNA was beads purified (1X ratio). RT was accomplished using random hexamers priming.

#### Generation of Gag-Pol amplicons

The cDNA of the 9 HIV-1 clinical samples and the HIV-1 control sample were used to generate amplicons. To remove excess primers, the resulting cDNA was beads purified (0.5X ratio) and eluted with 30µl nuclease-free water. Fifteen microliters of each sample were then used for PCR amplification using SuperFi DNA polymerase. To amplify ~2500 bp spanning entire Gag and part of Pol HIV-1 regions (HXB2 coordinates 524-3249, the following primers were used: GAG FW 5’CTC AAT AAA GCT TGC CTT GAG TGC and RT gene RV 5’ACT GTC CAT TTA TCA GGA TGG AG, and the following PCR program: initial denaturation for 3min at 98°C, followed by 40 cycles of denaturation for 20sec at 98°C, annealing for 30sec at 62°C and extension for 2.5min at 72°C, and final extension for 5min at 72°C. The amplicons were gel purified and their concentration was determined. The purified products were further used for library construction.

#### Generation of Gag amplicon with primer-ID from HIV9 sample

A primer specific to the entire Gag gene of HIV-1 (HXB2 position 2347) was designed with a 15 N-bases unique barcode followed by a linker sequence for subsequent PCR, Gag ID RT 5’TAC CCA TAC GAT GTT CCA GAT TAC GNN NNN NNN NNN NNN NAC TGT ATC ATC TGC TCC TG TRT CT. Based on the measured viral load and sample concentration, 4 µl (containing roughly 300,000 HIV-1 copies) were taken for reverse transcription reaction. Reverse transcription was performed using SuperScript IV RT with the following adjustments: (1) In order to maximize the primer annealing to the viral RNA, the sample was allowed to cool down gradually from 65°C to room temperature for 10 minutes before it was transferred to ice for 2min; And (2) The reaction was incubated for 30min at 55°C to increase the overall reaction yield. To remove excess primers, the resulting cDNA was beads purified (0.5X ratio) and eluted with 35µl nuclease-free water. To avoid loss of barcoded primers (“primer-ID”s) due to coverage drop at the ends of a read as a result of the NexteraXT tagmenation process (see "Library construction for Miseq"), the PCR forward primer was designed with a 60bp overhang so the barcode is far from the end of the read. The primers used for amplification were Gag ID FW 5’CTC AAT AAA GCT TGC CTT GAG TGC and Gag ID RV 5’AAG CGA GGA GCT GTT CAC TGC CAT CCT GGT CGA GCT ACC CAT ACG ATG TTC CAG ATT ACG. PCR amplification was accomplished using SuperFi DNA polymerase in a 50µl reaction with 33.5µl of the purified cDNA as input using the following conditions: initial denaturation for 3min at 98°C, followed by 40 cycles of denaturation for 20sec at 98°C, annealing for 30sec at 60°C and extension for 1min at 72°C, and final extension for 2min at 72°C. The Gag amplicon was gel purified and the concentration was determined. The purified product was further used for library construction.

#### Generation of Vif amplicon from HIV9 sample

One and a half microliters from clinical sample HIV9 were reverse transcribed using SuperScript IV RT and random hexamer priming. Five microliters of the purified RT reaction were used to set-up a PCR reaction with SuperFi DNA polymerase to amplify ~600 bp region spanning HIV-1 Vif gene using primers vif FW 5’AGG GAT TAT GGA AAA CAG ATG GCA GGT and vif RV 5’CTT AAG CTC CTC TAA AAG CTC TAG TG, and the following program: initial denaturation for 3min at 98°C, followed by 40 cycles of denaturation for 20sec at 98°C, annealing for 30sec at 60°C and extension for 30min at 72°C, and final extension for 5min at 72°C. The amplicon was gel purified and the concentration was determined. The purified product was further used for library construction.

### Generation of amplicons from RSV clinical samples

#### Clinical RSV samples

Nasopharyngeal samples of 25 patients hospitalized at Chaim Sheba Medical Center (Table S6) were collected into Virocult liquid viral transport medium (LVTM) (Medical Wire & Equipment Co, Wiltshire, United Kingdom) and stored at −70°C. Five hundred microliters of each sample were extracted and purified using easyMAG according to the manufacturer’s instructions. A primer specific to the glycoprotein protein G was designed with a 15 N-bases unique barcode followed by a linker sequence for subsequent PCR, RSV G RT 5’TAC CCA TAC GAT GTT CCA GAT TAC GNN NNN NNN NNN NNN NGC AAA TGC AAM CAT GTC CAA AA. Eight microliters of each sample were reverse transcribed as described in “Generation of Gag amplicon with primer-ID from HIV-1” section.

#### RSV control sample

In the absence of an RSV plasmid, we used human rhinovirus (RV) plasmid (a kind gift by Ann Palmenberg (University of Wisconsin-Madison, WI, USA) to generate a homogeneous control that was run on the same Nextseq run as the RSV samples, similar to the described above. In vitro transcribed RNA underwent RT using SuperScript IV with the following primer: RV14 5’ TAC GCA TAC GAT GTT CCA GAN NNN NNN NNN NNN NNN NNN NAT AAA CTC CTA CTT CTA CTC AAA TTA AGT GTC. PCR amplification using Q5 DNA polymerase with the following primers was performed: p3.26 FW 5’ TTA AAA CAG CGG ATG GGT ATC CCA C and p3.26 RV 5’ATG GTG AGC AAG GGC GAG GAG CTG TTC ACC GGG GTG GTG CTA CGC ATA CGA TGT TCC AGA.

#### Generation of a glycoprotein-fusion protein amplicon and polymerase amplicons

PCR amplification was accomplished using Q5 DNA polymerase in 50µl reactions with 15µl of the purified cDNA as input. The following conditions were used for the glycoprotein-fusion protein amplicon: initial denaturation for 3min at 98°C, followed by 40 cycles of denaturation for 20sec at 98°C, annealing for 30sec at 58°C and extension for 3.5min at 72°C, and final extension for 5min at 72°C, using the following primers: Extension FW 5’AAG CGA GGA GCT GTT CAC TGC CAT CCT GGT CGA GCT ACC CAT ACG ATG TTC CAG ATT ACG and RSV G and F RV 5’TGA CAG TAT TGT ACA CTC TTA. For the polymerase amplicon, the following conditions were used: initial denaturation for 3min at 98°C, followed by 40 cycles of denaturation for 20sec at 98°C, annealing for 30sec at 60°C and extension for 8min at 72°C, and final extension for 5min at 72°C, using the following primers: RSV L FW 5’GGA CAA AAT GGA TCC CAT TAT T and RSV L RV 5’GAA CAG TAC TTG CAY TTT CTT AC. The amplicons were beads purified and joint together at equal amounts. Concentration was determined, and the product was further used for NextSeq library construction.

### Generation of a UL54 amplicon from CMV clinical samples

Clinical DNA samples of recently infected patients (see Table S6) were obtained and purified as described previously (Weisblum, et al. 2017). Since CMV is a DNA virus, no reverse transcription step was needed. To generate a homogeneous control sample, the UL54 gene from TB40/E strain was cloned onto a pGEM-t plasmid as described previously (Weisblum, et al. 2017). The samples were diluted to 30,000 copies per PCR amplification reaction, which was set-up using the Q5 DNA polymerase. The primers used to amplify the UL54 gene were UL54 FW 5’TCA ACA GCA TTC GTG CGC CTT and UL54 RV 5’ATG TTT TTC AAC CCG TAT CTG AGC GGC, and the following PCR protocol was executed: initial denaturation for 3min at 98C, followed by 38 cycles of denaturation for 20sec at 98C, annealing for 20sec at 65C and extension for 3min at 72C, and final extension for 5min at 72C. The amplicons were beads purified and their concentrations were determined. The purified products were further used for MiSeq library construction with the following change, 0.875ng of DNA were used as input for tagmentation instead of 0.85ng.

### MiSeq/Nextseq Libraries construction

PCR fragmentation and indexing of samples for sequencing was performed using the Nextera XT DNA Library Prep Kit (Illumina, San Diego, CA, USA) with the following adjustments to the manufacturer instructions; (1) In order to get a short insert size of ~250bp, 0.85 ng of input DNA was used for tagmentation; (2) No neutralization of the tagmentation buffer was done, as described previously (Baym, et al. 2015); (3) For library amplification of the tagmented DNA, the Nextera XT DNA library prep PCR reagents were replaced with high-fidelity DNA polymerase reagents (the same DNA polymerase that was used for the amplicon generation). The PCR reaction (50µl total) was set as depicted. Directly to the tagmented DNA, index 1 (i5, illumina, 5µl), index 2 (i7, illumina, 5µl), buffer (10µl), high-fidelity DNA polymerase (0.5µl), dNTPs (10mM, 1µl) and nuclease-free water (8.5µl) were added; (4) Amplification was performed with annealing temperature set to 63°C instead of 55°C, as introduced previously (Baym, et al. 2015) and final extension for 2min; (5) Post-amplification clean-up was achieved using AMPure XP beads in a double size-selection manner (Bronner, et al. 2014), to remove both too large and too small fragments in order to maximize the fraction of fully overlapping read pairs. For the first size-selection, 32.5µl of beads (0.65X ratio) were added to bind the large fragments. These beads were separated and discarded. For the second-size selection, 10µl of beads (0.2X ratio) were added to the supernatant to allow binding of intermediate fragments, and the supernatant containing the small fragments was discarded. The intermediate fragments were eluted and their size was determined using a high-sensitivity DNA tape in Tapestation 4200 (Agilent, Santa Clara, CA, USA). A mean size of ~370bp, corresponding to the desired insert size of ~250bp, was achieved; And (6) Normalization and pooling was performed manually.

NextSeq: The longest NextSeq read length is 150bp, we hence selected for a shorter insert size of 270bp, compared to the desired 370bp insert size for the MiSeq platform. The first size selection of the post-NexteraXT amplification cleanup was performed using 42.5µl of AMPure XP beads (0.85X ratio) (Bronner, et al. 2014).

### AccuNGS Development

The AccuNGS protocol was evaluated using HIV-1 DNA plasmid (Peden, et al. 1991)). Our underlying assumption was that this DNA starting material is homogenous with respect to the theoretical error rate we calculated. This assumption was based on the fact that we used low-copy plasmids that were grown in *Escherichia coli*, and only a single colony was subsequently sequenced. The mutation rate of *E. coli* is in the order of 1×10^−10^ errors/base/replication (Jee, et al. 2016), and accordingly, error rates in the purified plasmids are expected to be much lower than the original expected protocol mean error of ~10^−5^ (Table S7). Table S1 summarizes the different differential sequencing that was performed during the Methods development, in an attempt to test the contribution of each one of the stages of the protocol to the error rate of the method (see Supplementary Text).

#### Preparation of plasmids

In order to maintain the plasmid stock as homogenous as possible, plasmids were transformed to a chemically competent bacteria cells [DH5alpha (BioLab, Israel) or TG1 [A kind gift by Itai Benhar (Tel Aviv University, Tel Aviv, Israel)]] using a standard heat-shock protocol. Based on the fact that *E. coli* doubling time is 20 minutes in average using rich growing medium (Sezonov, et al. 2007), a single colony was selected and grown to a maximum of 100 generations. Plasmids were column purified using HiYield™ Plasmid Mini Kit (RBC Bioscience, New Taipei City, Taiwan) and stored at −20°C until use.

#### Construction of baseline control amplicon

A baseline control amplicon (Table S1) was based on clonal amplification and sequencing of the pLAI.2 plasmid, which contains a full-length HIV-1_LAI_ proviral clone (Peden, et al. 1991) (obtained through the NIH AIDS Reagent Program, Division of AIDS, NIAID, NIH: pLAI.2 from Dr. Keith Peden, courtesy of the MRC AIDS Directed Program). The Integrase region of pLAI.2 was amplified using primers: KLV70 - 5’TTC RGG ATY AGA AGT AAA YAT AGT AAC AG and KLV84 - 5’TCC TGT ATG CAR ACC CCA ATA TG (Moscona, et al. 2017). PCR amplification was conducted using SuperFi DNA Polymerase in a 50µl reaction using 20-40 ng of the plasmid as input. Amplification in a thermal cycler was performed as follows: initial denaturation for 3min at 98°C, followed by 40 cycles of denaturation for 20sec at 98°C, annealing for 30sec at 60°C and extension for 1min at 72°C, and final extension for 2min at 72°C. In parallel, an alternative PCR reaction was up using Q5 DNA Polymerase. The Integrase amplicon was gel purified and concentration was determined. The purified product was further used for library construction.

#### Construction of AmpR and RpoB control amplicons

For generating the AmpR amplicon, the conserved *AmpR* gene was amplified from pLAI.2 plasmid using primers: AmpR FW - 5’AAA GTT CTG CTA TGT GGC GC and AmpR RV - 5’GGT CTG ACA GTT ACC AAT GC. PCR amplification was carried out as described above, except for extension duration of 30sec instead of 1min. Similarly, the conserved *RpoB* gene was amplified from the bacteria genome using the following primers: RpoB FW 5’ATG GTT TAC TCC TAT ACC GA and RpoB RV 5’GTG ATC CAG ATC GTT GGT G and the following PCR program: initial denaturation for 3min at 98°C, followed by 40 cycles of denaturation for 10sec at 98°C, annealing for 10sec at 60°C and extension for 4sec at 72°C, and final extension for 2min at 72°C. The AmpR and RpoB amplicons were gel purified and their concentration was determined. The purified product was further used for library construction.

#### Construction of alternative purification amplicons

The agarose gel purification step of the amplified integrase gene was replaced with other purification methods; (1) For the gel-free sample, the amplified integrase gene was purified using 25µl of AMPure XP beads (0.5X ratio), and (2) For the ExoSap sample, 10µl of the amplified integrase gene were mixed with 4µl of ExoSAP-IT™ PCR Product Cleanup Reagent (Thermo Fisher Scientific) and incubated according to the manufacturer’s instructions. No other changes in the generation of amplicon protocol were made.

#### Construction of a PCR-free control amplicon

For the PCR-free sample, 10µg of pLAI.2 plasmid was digested using the restriction enzymes: NheI, StuI and XcmI (NEB) according to the manufacturer’s instructions. A ~1500bp fragment containing the integrase region was gel purified and concentration was determined by Qubit. The purified product was further used for library construction.

#### Construction of RNA control amplicons

We used a plasmid containing the full cDNA of Coxsackie virus B3 (CVB3) under a T7 promoter that was a kind gift from Marco Vignuzzi (Institut Pasteur, Paris, France). Ten micrograms of this plasmid were linearized using SalI (NEB), beads purified (0.5X ratio) and then *in-vitro* transcribed using T7 RNA polymerase (NEB) according to the manufacturer’s instructions. The transcribed RNA was bead purified (0.5X ratio) and reverse transcribed with random hexamers using SuperScript III RT. Four microliters of the reverse transcription reaction were used as template for a PCR reaction using primers: CVB FW 5’GGA GAG AAG GTC AAC TCT ATG GAA GC and CVB RV 5’TAC CAC CCT GTA GTT CCC CA, which amplify a ~1500bp fragment within the CVB genome. PCR reaction (50µl total) was set and amplified using SuperFi DNA polymerase as follows: initial denaturation for 3min at 98°C, followed by 40 cycles of denaturation for 20sec at 98°C, annealing for 30sec at 60°C and extension for 15sec at 72°C, and final extension for 2min at 72°C. The CVB amplicon was gel purified and the concentration measured. The purified product was further used for library construction.

#### Alternative library purification methods

For the AMPure XP beads-free sample, post-amplification clean-up by double size-selection was replaced with an agarose gel purification of a ~370bp fragment, with no other changes in the library construction protocol.

#### Alternative tagmentation sample

For the alternative tagmentation sample, a 250bp amplicon within the integrase region was designed, using specific primers with an overhang corresponding to the sequence inserted during the tagmentation step of the NexteraXT DNA library prep kit, NexteraXT free FW 5’TCG TCG GCA GCG TCA GAT GTG TAT AAG AGA CAG ACT TGT CCA TGC ATG GCT TCT C and NexteraXT free RV 5’GTC TCG TGG GCT CGG AGA TGT GTA TAA GAG ACA GTC TAT CTG GCA TGG GTA CCA GCA. PCR reaction was set up using SuperFi DNA polymerase and carried out as follows: initial denaturation for 3min at 98°C, followed by 40 cycles of denaturation for 20sec at 98°C, annealing for 30sec at 62°C and extension for 15sec at 72°C, and final denaturation for 2min at 72°C. The PCR product was gel purified and the concentration was measured by Qubit. The purified product was indexed by a succeeding PCR amplification using primers corresponding to i5 and i7 NexteraXT primers (IDT) as mentioned previously (Baym, et al. 2015) at a final concentration of 1uM. The PCR reaction was set up using SuperFi DNA polymerase and amplified as detailed: initial denaturation for 3min at 98°C, followed by 12 cycles of denaturation for 20sec at 98°C, annealing for 30sec at 63°C and extension for 30sec at 72°C, and final extension for 2min at 72C. Size selection was achieved by gel purification of ~370bp fragments.

#### Sequencing

Sequencing of all synthetic samples, the HIV-1, and CMV samples was performed on the Illumina MiSeq platform using MiSeq Reagent Kit v2 (500-cycles, equal to 250×2 paired-end reads) (Illumina). Sequencing of the RSV-1 samples and a dedicated synthetic sample was performed on the Illumina NextSeq 500 platform using NextSeq 500/550 High Output Kit (300-cycles, equal to 150×2 paired-end reads) (Illumina).

#### Construction of synthetically mixed populations

A pLAI.2 plasmid was mixed with a pNL4.3 plasmid [a kind gift from Eran Bacharach (Tel Aviv University, Tel Aviv, Israel)] using serial dilutions to achieve the following ratios: 1:100, 1:500, 1:1,000, 1:5,000 and 1:10,000 with pLAI.2 held as the major strain. Concentrations were measured and each dilution was generated independently to obtain three technical replicas for each ratio. The plasmids mixture was further used for library construction.

### Barcode serial dilution test

The pLAI.2 plasmid was used to generate an RNA pool. Five micrograms of this plasmid were linearized using SalI (NEB) and beads purified (0.5X ratio). T7 polymerase promotor was added to the linearized plasmid using T7 extension FW 5’TAA TAC GAC TCA CTA TAG CTG GGA GCT CTC TGG CTA AC and the RV 5’GAG ACT CCC TGA CCC AGA TGC C in a PCR reaction using Q5 DNA polymerase with the following program: initial denaturation for 3min at 98°C, followed by 40 cycles of denaturation for 10sec at 98°C, annealing for 10sec at 65°C and extension for 3min at 72°C, and final extension for 5min at 72°C. Four microliters of the reaction was in-vitro transcribed using T7 RNA polymerase according to the manufacturer’s instructions. The transcribed RNA was beads purified (0.5X ratio). The purified RNA was serially diluted and for each dilution two reactions were set-up: a primer-ID reaction (as described in the section “Generation of Gag amplicon with primer-ID from HIV-1”) and a random hexamer based RT reaction (as described in the section “Construction of RNA control amplicons”). In order to compare these reactions, for the PCR amplification of the random hexamer based RT reaction, we used the following primers: GAG FW 5’CTC AAT AAA GCT TGC CTT GAG TGC and RTgene RV 5’ACT GTA TCA TCT GCT CCT GTA TCT corresponding to the primer-ID reaction primers without a barcode. The same PCR program was used for both reactions. The PCR reactions were gel purified and concentration was measured.

### Reads processing and base calling

The paired-end reads from each control library were aligned against the reference sequence of that control using an in-house script that relies on BLAST command-line tool (Altschul, et al. 1990; Altschul, et al. 1997; Camacho, et al. 2009). The paired-end reads from the clinical samples were aligned against: HIV-1 subtype B HXB2 reference sequence (GenBank accession number K03455.1), RSV reference sample (GenBank accession number U39661), CMV reference sample Merlin (GenBank accession number NC_006273), and then realigned against the consensus sequence obtained for each sample. Bases were called using an in-house script only if the forward and reverse reads agreed and their average Q-score was above an input threshold (30 or 38). At each position, for each alternative base, we calculate mutation frequencies by dividing the number of reads bearing the mutation by loci coverage. Positions were retained for analysis only if sequenced to a depth of at least 100,000 reads. In order to analyze the errors in the sequencing process we used Python 3.7.3 (Anaconda distribution) with the following packages: pandas 0.25.1 (McKinney 2010), matplotlib 3.1.0 (Caswell, et al. 2019), seaborn 0.9.0 (Waskom, et al. 2018), numpy 1.16.3 (Oliphant 2006; Walt, et al. 2011) and scipy 1.2.1 (Jones, et al. 2016). Distributions of errors on control plasmids were compared using two-tailed t-test or two-tailed Mann-Whitney U test.

### Variant calling

In order to facilitate discrimination of true variants from AccuNGS process artifacts, we created a variant caller based on two principles: (i) positions that exhibit relatively high level of error on a control sample are error-prone for the clinical sample as well; and (ii) process errors on a control sample follow a gamma distribution. A gamma distribution was fitted for each mutation type in the control sequence. In order to detect and remove outliers from the fitting process we used the “three-sigma-rule”, and positions that showed error higher than three standard deviations from the mean of the fitted distribution were removed. For these rare loci a base was called only if the mutation was more prevalent in the sample by an order of magnitude. For G>A transition mutations, four distinct gamma distributions were fitted, corresponding for all four G>A combinations with preceding nucleotide. Accordingly, for C>T transition mutations four gamma distributions were fitted as well, on the four C>T reverse complement mutations of the G>A mutations. For establishing Figures 2-5, variants were called on the input sample only if a mutation was in the extreme 1% of the relevant gamma distribution fitted using the corresponding control.

### Serial dilutions analysis

All dilutions were mapped against the pLAI.2 reference sequence using the default parameters and Q30 as the minimal base quality threshold. Positions with insufficient coverage, defined as having less than 5 times the inverse of the frequency of the minor strain were filtered out, as well as positions with minor variant frequency of above 0.5% in the homogeneous control sample.

### Diversity calculation

Transition nucleotide diversity π was calculated per sample using positions with at least 5,000x coverage, using the formulas described in (Zhao and Illingworth 2019), but excluding transversion variants. Sites whose variant frequencies weren’t statistically different than the background control were considered as variants of count = 0 for the calculation.

### Haplotypes inference

To infer potential haplotypes, we used a two-step process, illustrated in Figure 6. First we identified all pairs of non-consensus variants (the most common minor variant at each site) that were statistically enriched when present on the same reads. Next we attempted to "concatenate" multiple pairs into a longer stretch based on a shared mutation present in two different pairs of variants. In order to find statistically enriched pairs, we considered all sites that may be linked on the same reads (up to 250 bases, which is the maximal length of an Illumina read). For each pair of loci we created a contingency table for the appearance of each variant alone, the two variants together and no variant at all. We then used a one-tailed Fisher exact test to obtain a p-value for the pair, and considered only p-values lower than 10^−15^, to account of multiple testing. From this contingency table we also extracted the frequency at which the two variants co-occur. We repeated the process for all possible pairs of loci. This resulted in many short haplotype stretches of 250 bases spanning two loci each. We then performed "concatenation" of pairs of loci that had (1) at least one shared position and (2) a similar frequency of co-occurrence, defined here as up to an order of magnitude in difference. Such concatenated loci formed a longer stretch and its frequency was calculated as the mean frequency of its components, i.e., the average frequency of all individual pairs added to this stretch so far. For each sample, we iteratively attempted to concatenate all pairs of loci, starting from the highest frequency pair to the least common pair, until no pairs could further merge.

**Figure 6.**
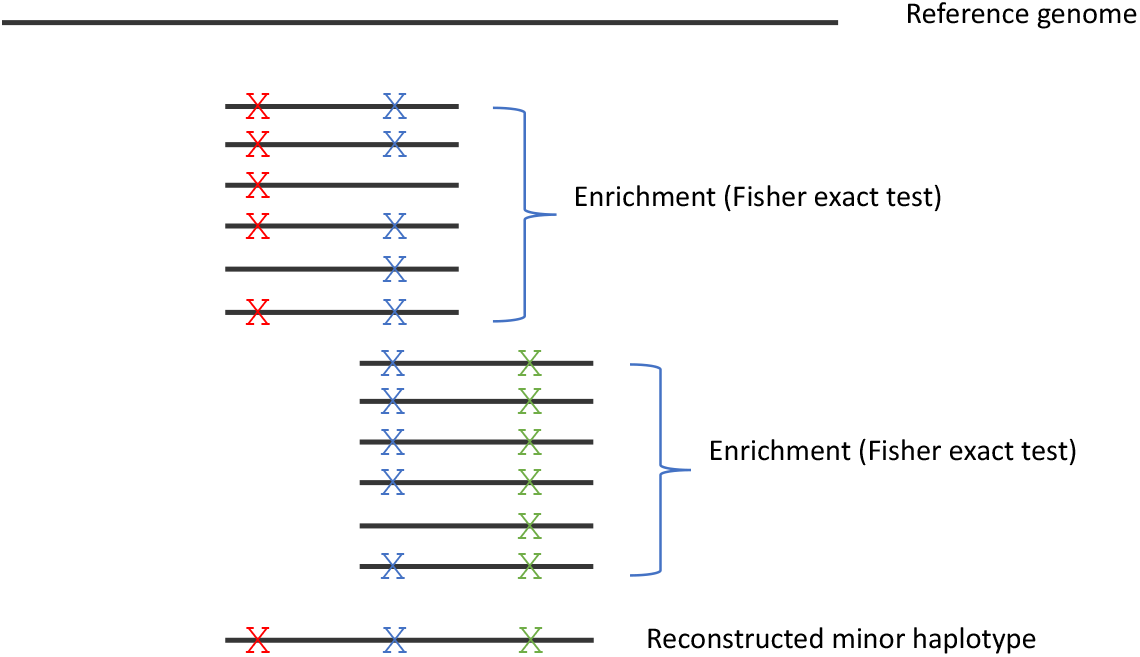
Illustration of method for haplotype reconstruction. The method searches for enrichment of pairs of mutations on the same read, and concatenation of enriched reads that share a mutation into a reconstructed minor haplotype. Notably, the concatenation approach is suitable for populations with limited diversity, as is the case in acute infections; in highly diverse populations, many haplotypes may share the “blue” mutation illustrated in the figure.

### CODE AVAILABILITY

We have developed the following computational resources that complement the AccuNGS sequencing protocol:

a. Base coverage calculator. AccuNGS relies on overlapping read pairs and high Q-scores for both reads of a pair. The calculator receives as input the length of the target regions and the desired coverage, and outputs the recommended number of reads required for sequencing each sample.
b. Computational pipeline for computing the number of unique RNA molecules sequenced, based on primer-ID barcodes (see Supplementary Text).
c. Computational pipeline for base-calling and inferring site by site base frequencies.
d. Computational pipeline for inferring haplotypes.

All resources are freely available at https://github.com/SternLabTAU/AccuNGS.

### ACCESSION NUMBERS

The datasets generated and reported in this study were deposited in the Sequencing Read Archive (SRA, available at https://www.ncbi.nlm.nih.gov/sra), under BioProject PRJNA476431. Frequencies of mutations following base calling will be available in Zenodo (https://zenodo.org/) upon acceptance.

## ACKNOWLEDGEMENTS

The authors would like to thank Oded Kushnir, Danielle Miller and Yiska Weisblum for valuable support, and for Drs. Neta Zuckerman, Tzachi Hagai and Shaul Pollak for critical reading of the manuscript and helpful discussions.

