## Supplementary Material for "Accurate *in vivo* population sequencing uncovers drivers of within-host genetic diversity in viruses"

**SUPPLEMENTARY TEXT 1 - The development and validation of the AccuNGS sequencing approach**

**AccuNGS error landscape at the DNA level.**

We compared the error rate of the baseline AccuNGS protocol (Table S3) to the results of the typical NGS protocol used when less emphasis is put on the fidelity of the process. Reassuringly, AccuNGS showed a significant improvement of one to two orders of magnitude over the standard sequencing protocol for most types of mutation (Fig. S1). For example, the mean A>G error rate went down from $2.5x10^{-3}$ down to around $6.21x10^{-5}$. While this improvement was large, the resulting error rate was still three times higher than the theoretical error rate we had expected of ~$2x10^{-5}$. We performed a set of sequencing trials, where at each trial we tested if removing or changing a specific stage of the protocol alleviates some of the observed errors and improves the fidelity of AccuNGS (Table S1). We succeeded in ruling out that most errors are derived from any of the library preparation steps including (i) PCR, (ii) gel extraction, (iii) DNA size selection (particularly UV light exposure), (iv) tagmentation during the NexteraXT DNA library preparation kit (Illumina), (v) alternative PCR cleanup procedures (magnetic beads extraction and using the Exosap cleanup reagent), (vi) the bacteria used to grow the plasmid, (vii) the particular sequence of the plasmid, (viii) the sequencing machine (Table S2). Notably, in some samples we observed the potential effects of oxidation and/or deamination, the former consistent with elevated G>T and C>A in a particular context (Fig. S1C,D). Following these observations, we incorporated context specific errors into our bioinformatics variant caller.

We next set out to test whether the sequencing machine itself was responsible for the observed errors. We applied a very stringent quality filtering of Q38 on one of our very high coverage samples (*AmprR*, Table S1). We expected that this filtering would improve the results by the difference between twice Q30 (Q60, error probability of $1x10^{-6}$) and twice Q38 (Q76, error probability of ~$2x10^{-8}$). This difference translates to an improvement which is far below our observed error rate and hence we did not expect to see any improvement. Surprisingly, we observed a significant reduction in the rates of errors for A:T>G:C miscalls, and a modest yet significant reduction for C:G>T:A miscalls (Fig. S1B, Table S3). We hence concluded that the assumption that the Q-scores of overlapping reads are independent (Zhang, et al. 2014; Edgar and Flyvbjerg 2015) is an incorrect assumption, and that the sequencer itself is likely the major source of errors in AccuNGS.

Finally, we tested the error profile of AccuNGS when starting off with RNA, where an additional reverse transcription step is incorporated, potentially introducing some more errors. Indeed, our results showed slightly elevated error rates when the protocol was tested on *in vitro* transcribed RNA (Methods).

**SUPLLEMENTARY TEXT 2 - Quantifying the number of templates sequenced**

**Quantifying the amount of templates.** During the calibration of the AccuNGS protocol, we began by testing whether it would be possible to quantify the number of templates sequenced. To this end, we used uniquely barcoded primers during the reverse transcription process to uniquely tag each viral genome that underwent reverse transcription in order to quantify the number of sequenced templates (Kivioja, et al. 2011). This method is also known as the "primer ID" method (Jabara, et al. 2011), although here it was not used for correcting process errors, because error correction is not required when the process fidelity is very high as we report for AccuNGS. However, during the course of calibration of this approach, we noted that the mere addition of a primer-ID led to much lower yield of DNA when testing on *in vitro* transcribed RNA at different dilutions (Fig. S2), as compared to the use of random hexamer without a primer-ID.

We next split one of our samples (9) into two vials. Given the viral load of this sample we estimated that each vial contained ~300,000 viral genome copies. We sequenced both vials: one was sequenced using a 15-nt random primer-ID at the reverse-transcriptase stage and one was sequenced using random hexamers. Following the reverse transcription stage the subsequent AccuNGS stages are identical. This allowed us to count 15,880 unique primer-IDs in the first vial (see paragraph below), which is about 5% of the original number of copies. Since we observed that the use of random hexamers leads to more DNA product than the use of primer-ID, we hence conclude that we sequence at least 5% (and probably more) of the number of original templates using our AccuNGS approach based on random hexamers.

**Counting primer-IDs.** A common problem when relying on barcodes (primer-IDs) in the sequence is that the barcodes themselves are not error-prone and accumulate sequencing errors as well. When the number of sequenced templates is much lower than the sequencing depth, this may manifest as some primer IDs that are observed only a few times, and may be at a short "edit distance" from their source barcode. Such barcodes are often named "offsprings". Previous study has mapped the relationship between the most abundant barcode and the reliable depth a barcode is required to be sequenced in order to be considered as non-offspring (Zhou, et al. 2015). Notably, this relationship was inferred using an experimental protocol that is more error-prone than AccuNGS. Currently, no standard exists for analyzing the primer IDs and downstream analysis. Zhou et al. provided a computational script that identifies primer ID by finding sequenced reads that has a perfect match to the known flanking sequences of the degenerate barcoded region. However, as sequencing errors may perturb the flanking regions this may result in incomplete barcode recovery.

We therefore developed our own primer ID recovery algorithm using a local alignment of the immediate flanking regions of the sequenced reads. The number of unique primer IDs identified is therefore an upper bound on the number of actually-sequenced templates, which in our sample was 22,971.

**Offspring analysis.** As most sequenced unique barcodes were sequenced once or twice and suspected to be offsprings of some paternal barcode, we tested the Hamming distance between all pairs of barcodes. Our analysis revealed that 15,880 barcodes had their closest barcode at a hamming distance of at least 3, suggesting that in AccuNGS most sequenced barcodes are real, and low barcode abundance may occur due to overall lower coverage and not due to sequencing errors of a more abundant barcode. We therefore conclude that we sequenced at least 15,880 different viruses.

**Contamination across samples**. Following our haplotype reconstruction procedure, we initially noted the putative presence of an additional very low frequency haplotype in many of the HIV samples, and in some of the RSV samples. When examining these haplotypes, it turned out that they were often identical to a consensus sequence of another sample. This led us to suspect contamination may have occurred during (a) one of the stages of the library preparation, or (b) during sequencing, or (c) during de-multiplexing, when each read is assigned to a sample. We were able to mostly rule out the latter, by testing the hamming distance of the barcodes of the contaminated reads to the expected barcode of the sample. Notably, we took utmost care to avoid contamination during library preparation, including the fact that when samples were run together on a gel, we added an empty lane between each pair of samples, and used a different knife to cut out each band. Nevertheless, we cannot rule out that minute contamination occurred at any one of the three stages above. We thus flagged all contaminated reads and removed them from all subsequent analyses.

**SUPPLEMENTARY FIGURES**

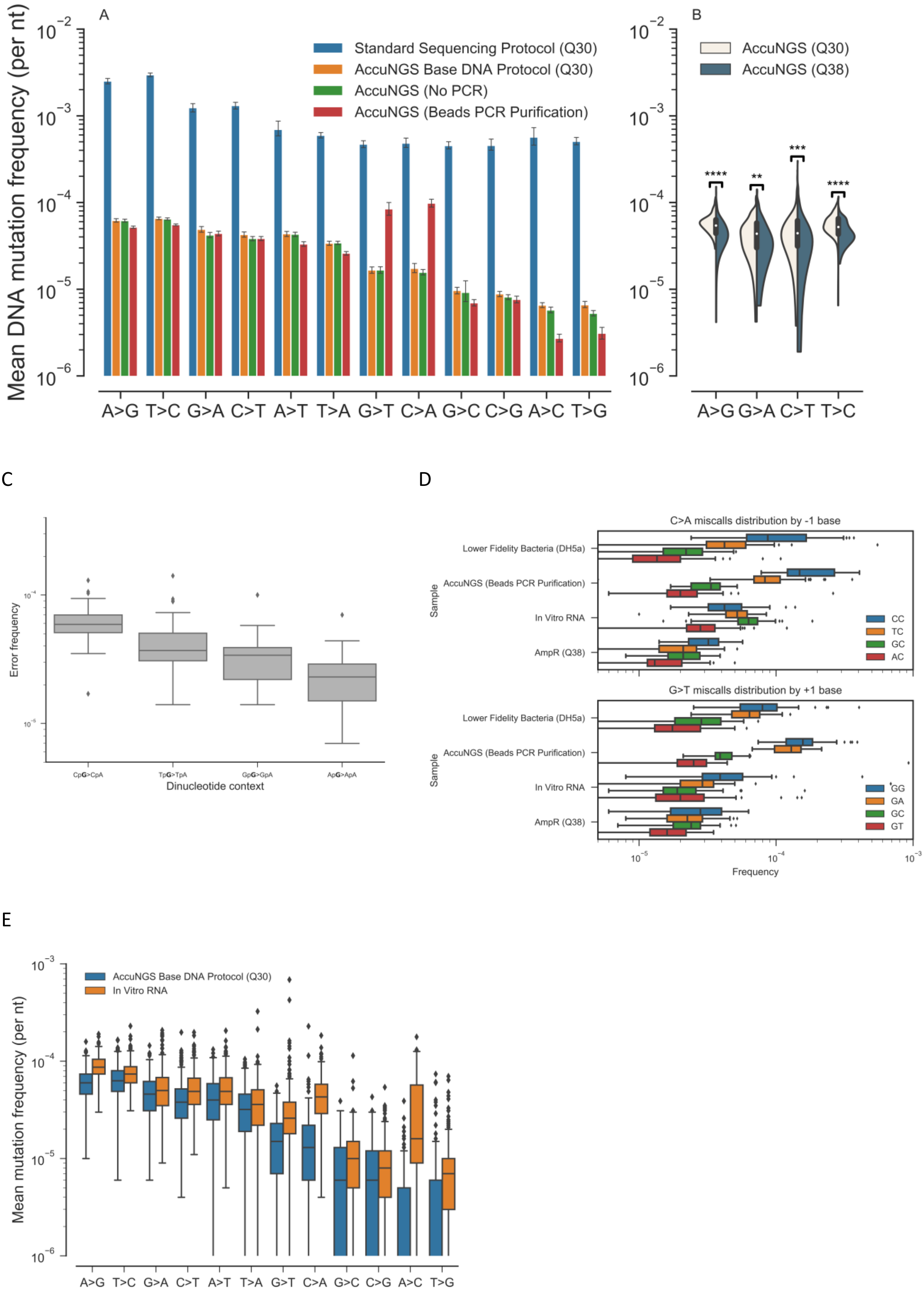

**Figure S1. Mean background error rates of different sequencing protocols at the DNA level.** (A) AccuNGS dramatically reduces errors present in standard sequencing protocols by almost two orders of magnitude. For standard sequencing control, a standard homogeneous pLAI.2 control was taken from (Moscona, et al. 2017), and mutations were called without accounting for overlapping paired reads, while considering positions to analysis only if sequenced to at least 2,000x depth. PCR errors in AccuNGS are negligible (in average) based on the comparison of a PCR and PCR-free sample. Higher rates of G>T and C>A are likely indicative of oxidative stress. Error bars represent 95% confidence intervals around estimated mean values using 1,000 bootstrap repeats. (B) The effect of increasing the Q-score filtering threshold on AccuNGS error rates, presented for each type of transition error. A>G and T>C transitions show the most dramatic effect when increasing the Q-score filtering threshold. (C) Effect of sequence context on G>A error frequencies potentially associated with deamination. Data shown on the *AmpR* control sample (Table S1) with Q38 filtering. (D) Distributions of process errors potentially associated with oxidative damage (G:C>T:A). Notably in the *In Vitro* RNA sample, the C>A errors pattern is different from the pattern observed in other samples due to the single-stranded origin of this sample. (E) Error rates of the DNA versus RNA control samples. Comparison of error distributions reveals that RT most often does not introduce a dramatically high level of error. Boxplots of errors per type of base changes are shown. Raw read bases were filtered when their average Q-score was less than Q30. **$p<0.01$; ***$p<0.001$; ****$p<0.0001$

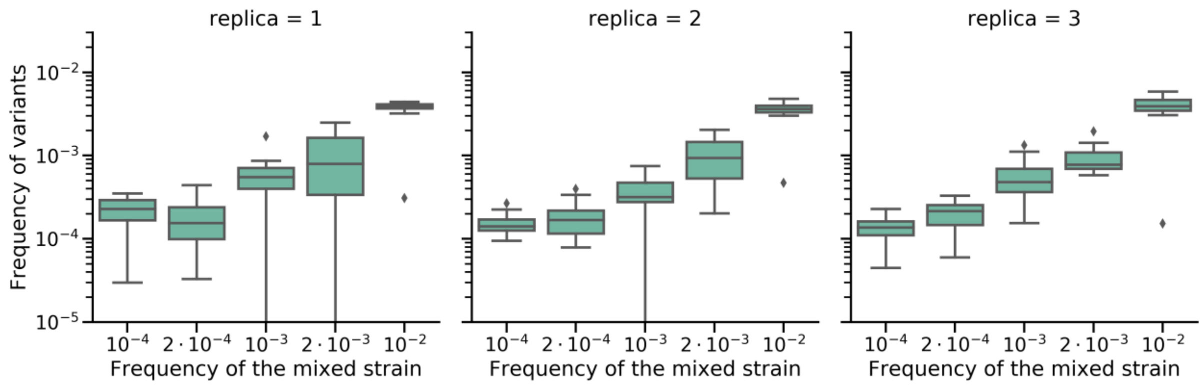


**Figure S2. Variant frequencies of all minor alleles in three serial dilution experiments of the synthetically created populations.** The x-axis represents the expected frequency of the infrequent plasmid, based on the dilution performed. The y-axis represents the frequency of variants that are present only on the infrequent plasmid. For all dilutions performed, the variant frequencies of the infrequent plasmid were extremely close to their expected frequency based on the dilution used.

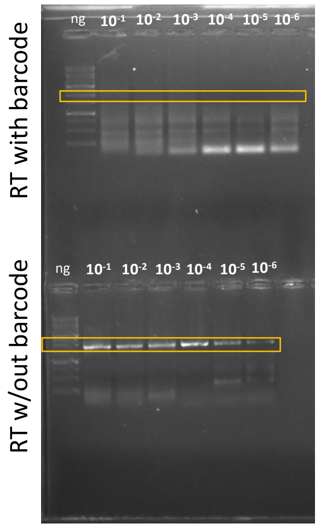


| ng | Estimated number of copies |
|---|---|
| 0.1 | $3.8x10^{7}$ |
| 0.01 | $3.8x10^{6}$ |
| 0.001 | $3.8x10^{5}$ |
| 0.0001 | $3.8x10^{4}$ |
| 0.00001 | $3.8x10^{3}$ |
| 0.000001 | $3.8x10^{2}$ |

**Fig. S3**. PCR results of serially diluted samples run with RT that includes a barcode (top) compared to RT without a barcode (bottom). The row corresponding to the estimated produced size (1935 bp with a barcode and 1860 bp without a barcode) is boxed. Estimated number of templates following dilution is shown on the right.

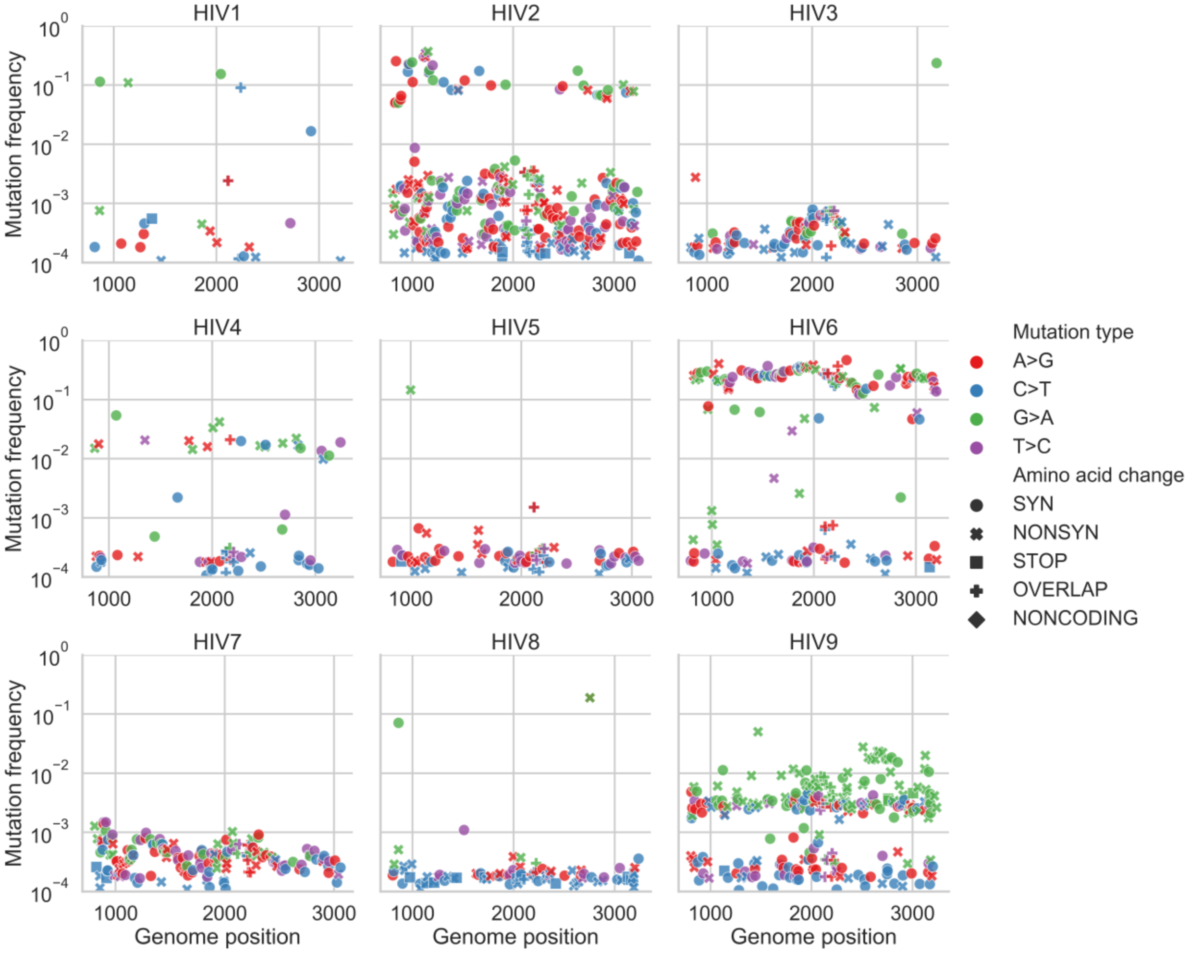
HIV1
HIV2
HIV3
HIV4
HIV5
HIV6
HIV7
HIV8
HIV9
Mutation frequency
Genome position
10^0
10^-1
10^-2
10^-3
10^-4
1000
2000
3000
Mutation type
A>G
C>T
G>A
T>C
Amino acid change
SYN
NONSYN
STOP
OVERLAP
NONCODING

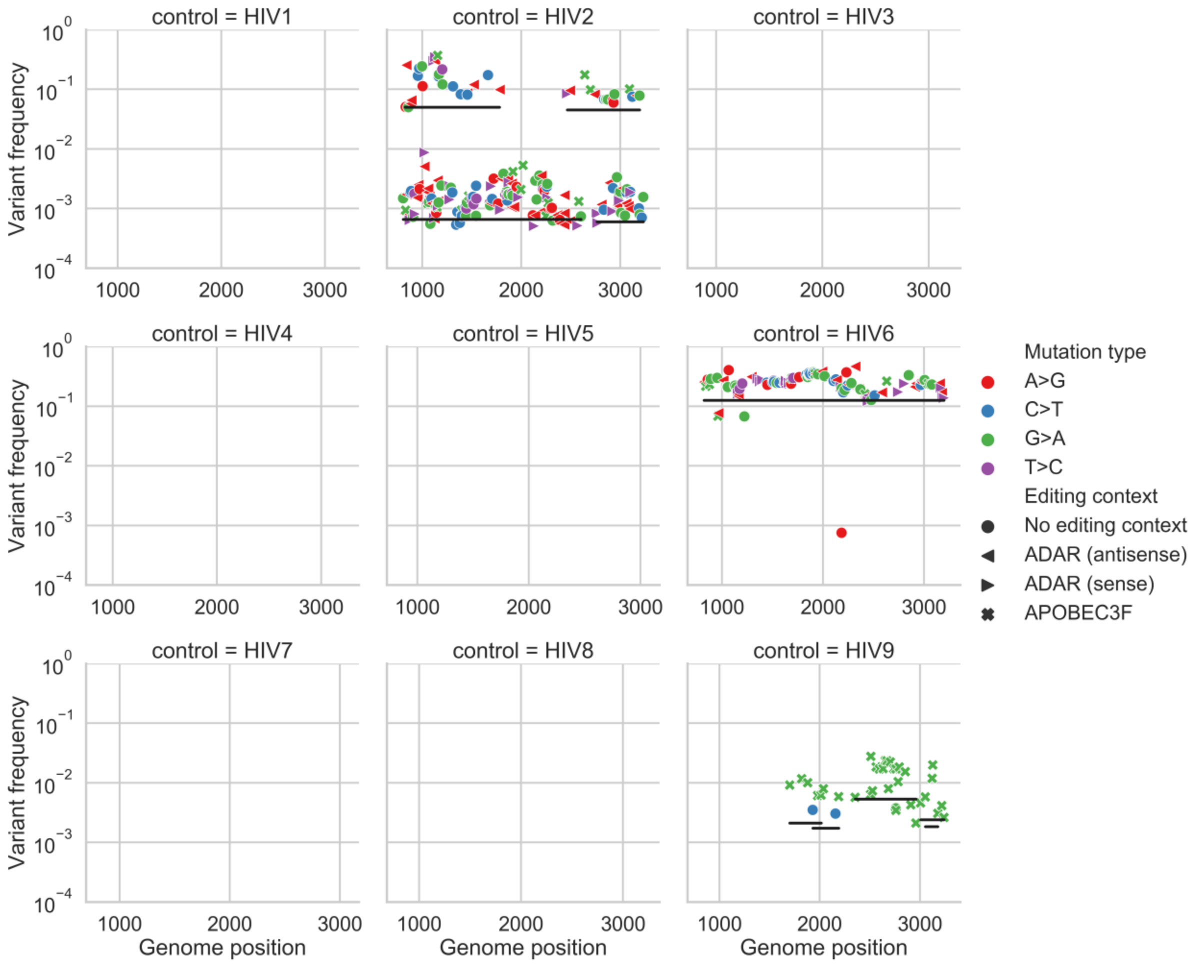


**Fig S4. Variant frequencies of HIV samples.** (A) Shown are transition variant frequencies along the sequenced gag-pol region of HIV, (B) Inferred haplotypes across all HIV samples. Details as in figures 3 and 4 of the main text respectively.

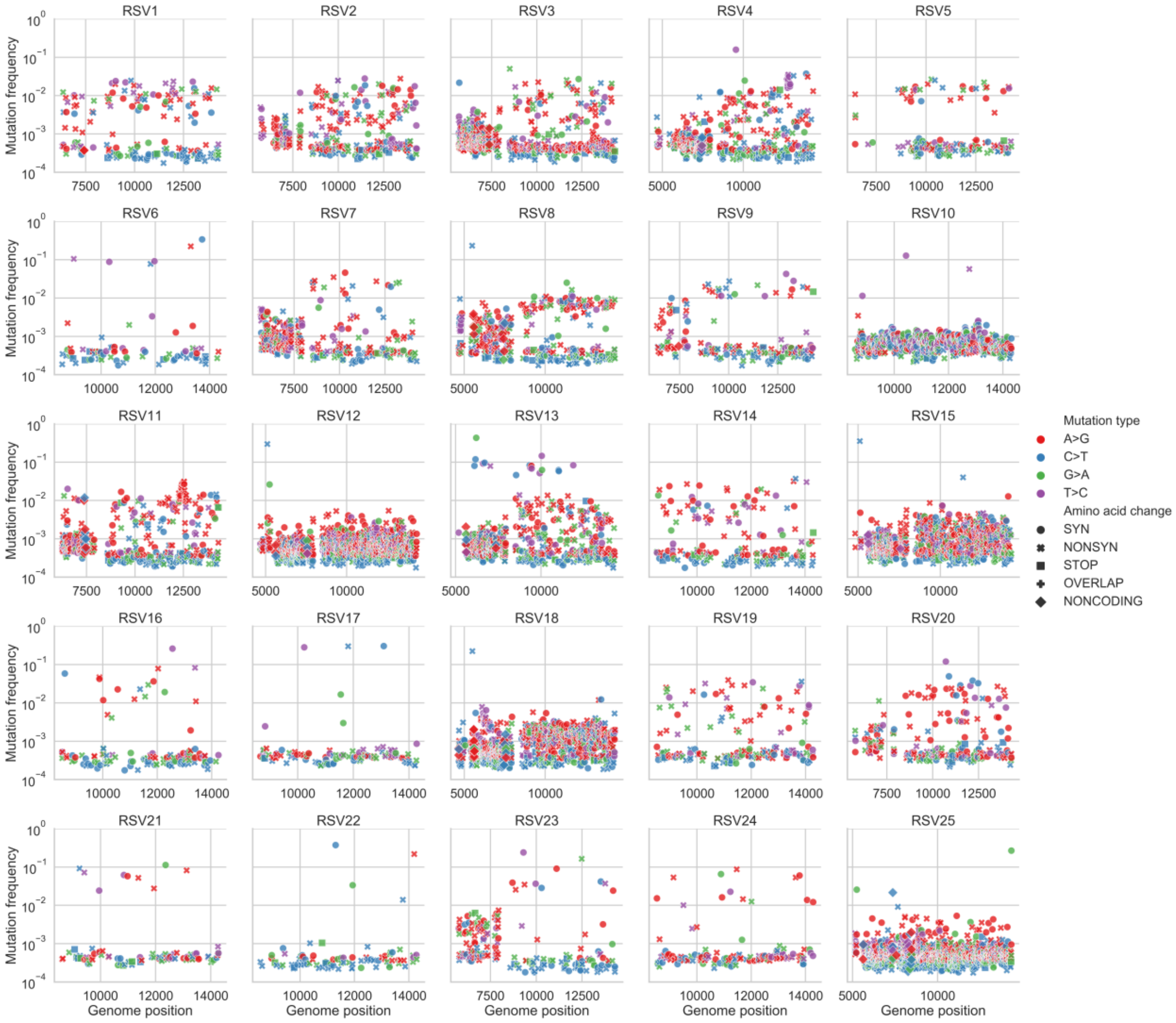
RSV1
RSV2
RSV3
RSV4
RSV5
RSV6
RSV7
RSV8
RSV9
RSV10
RSV11
RSV12
RSV13
RSV14
RSV15
RSV16
RSV17
RSV18
RSV19
RSV20
RSV21
RSV22
RSV23
RSV24
RSV25
Mutation frequency
Genome position
Mutation type
A>G
C>T
G>A
T>C
Amino acid change
SYN
NONSYN
STOP
OVERLAP
NONCODING

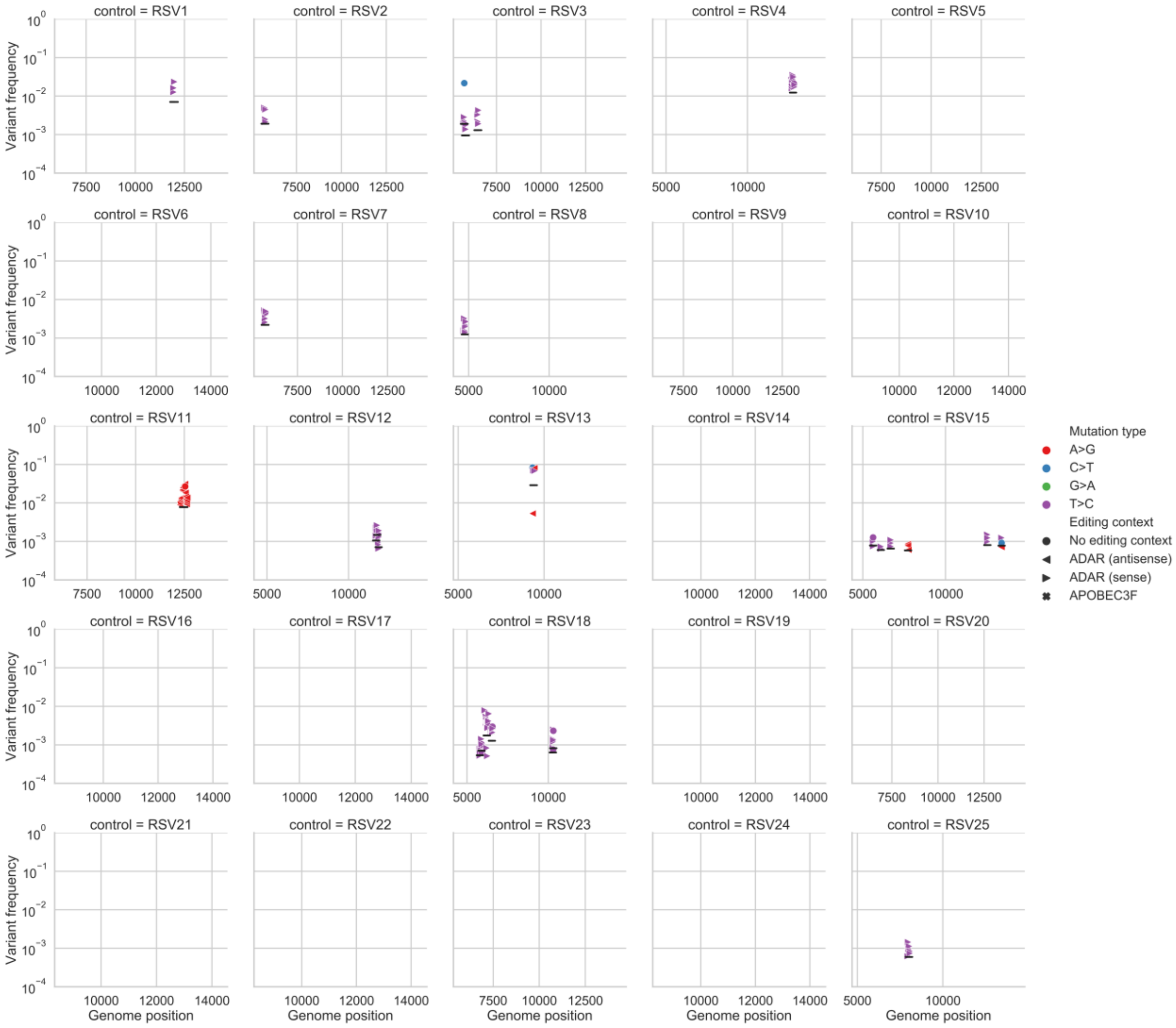


**Fig S5. Variant frequencies of RSV samples.** (A) Shown are transition variant frequencies along the sequenced regions of RSV. (B) Inferred haplotypes across all RSV samples. Details as in figures 3 and 4 of the main text respectively.

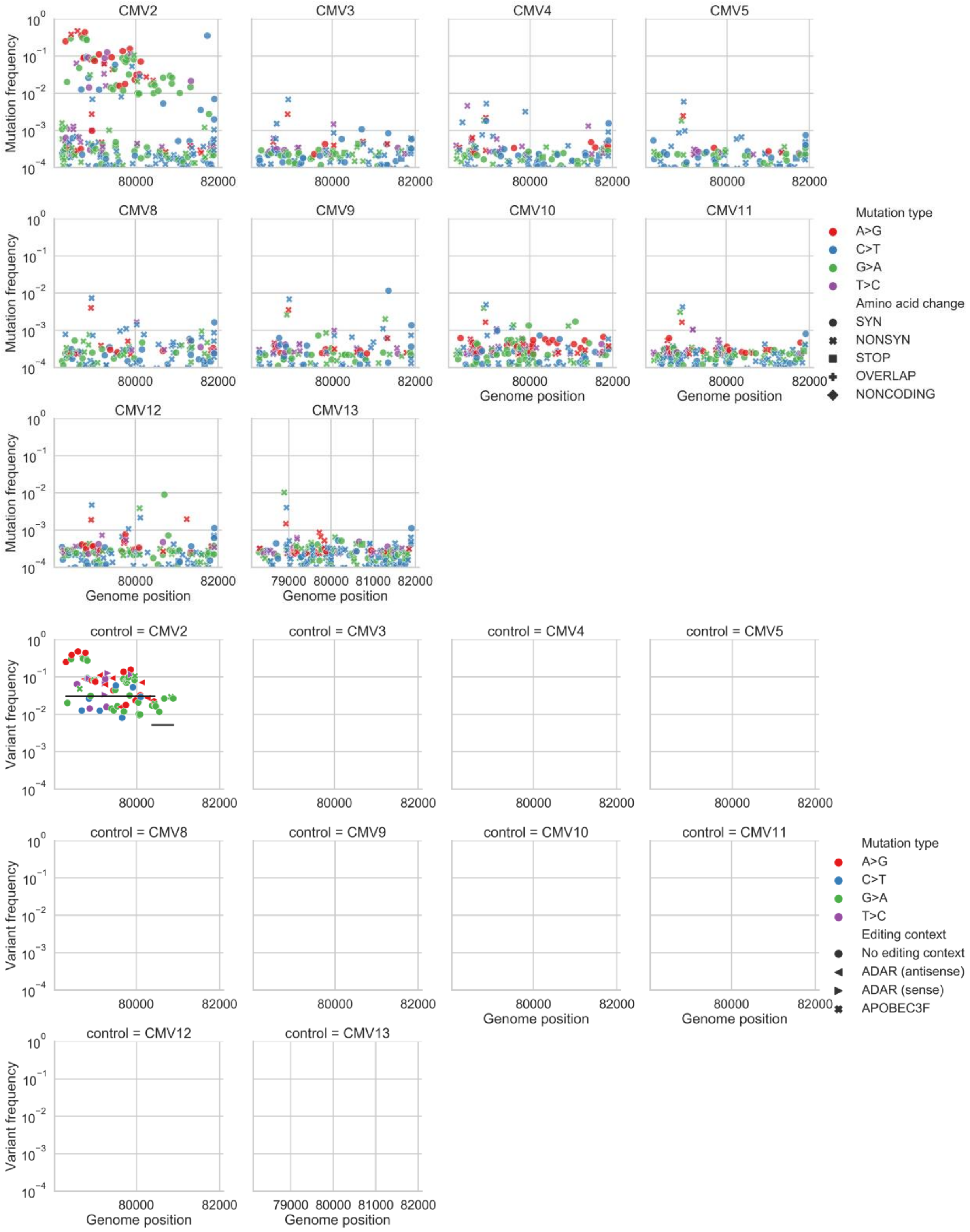


**Fig S6. Variant frequencies of CMV samples.** (A) Shown are transition variant frequencies along the sequenced regions of CMV. (B) Inferred haplotypes across all CMV samples. Details as in figures 3 and 4 of the main text respectively.

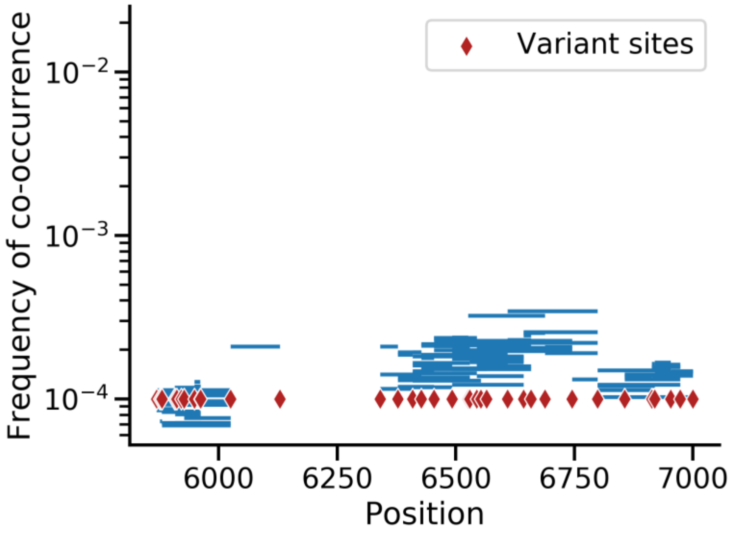


**Figure S7. Verification of haplotype reconstruction method on synthetically created populations.** Variants on the 1:10,000 mixed sample identified as linked to another variant on the same haplotype are shown as blue lines. All positions that differed between the two samples were found to be linked to each other in two large stretches that were distant enough to stay distinct. The only exception is one pair of variants that was not linked since the intermediate genomic region was identical between the two synthetically mixed strains. The frequencies of the constructed haplotypes approached the level of their dilution.

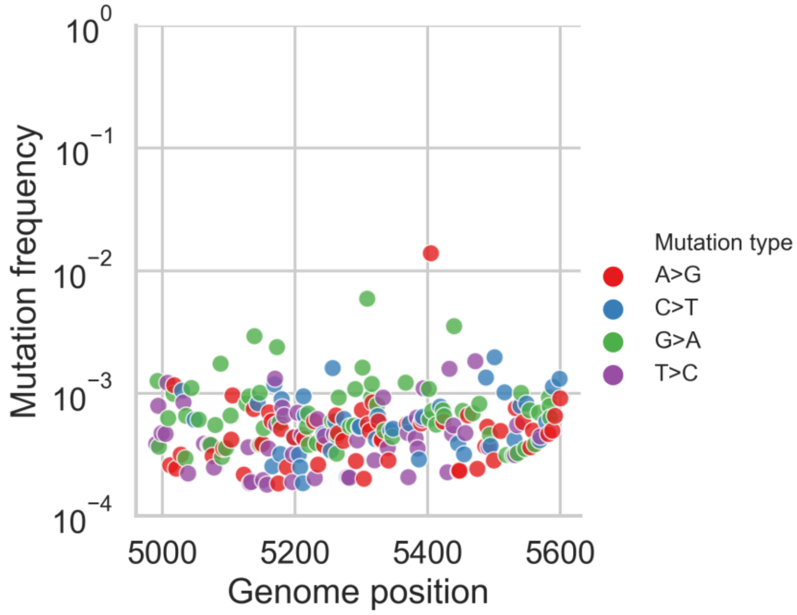


**Figure S8. Vif sequencing of HIV sample 6 reveals excess of G>A mutations in the context of APOBEC3D/F/H.**

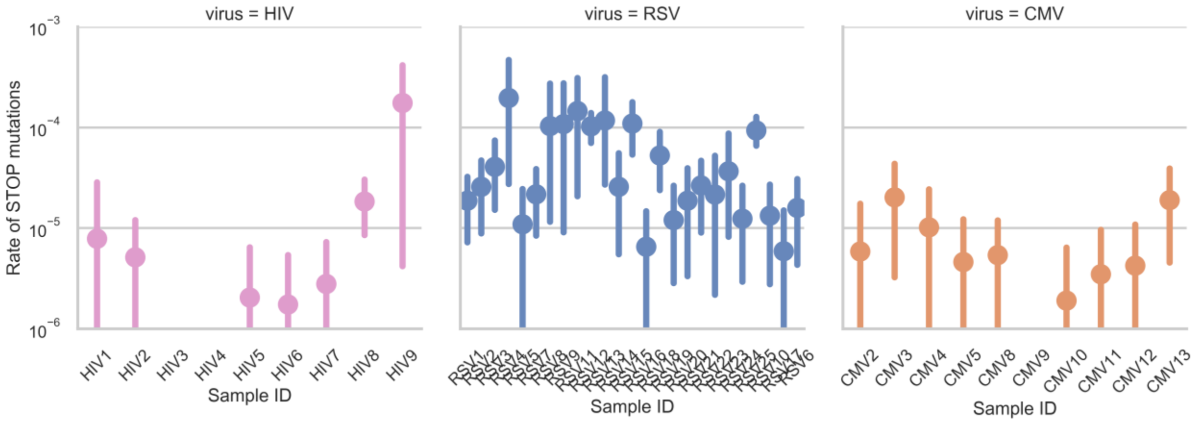


**Figure S9. Rates of stop mutations for all samples in this study.** The rate of a stop mutation is calculated as the number of reads bearing a premature stop codon divided by the total number of reads sequenced at loci covering potential sites that allow for premature stop codons. Bars represent 95% confidence intervals based on 1,000 bootstraps of these sites.

**SUPPLEMENTARY TABLES**

**Table S1. Differential DNA control samples sequenced in this study.** "." in a cell indicates a condition that is similar to the baseline. All samples besides the NextSeq sample were sequenced on the Illumina MiSeq sequencing platform.

| Sample name | PCR enzyme | PCR cycles | PCR purification method | Library purification method | Cloning bacteria | Tagmentation method | Target region |
|---|---|---|---|---|---|---|---|
| **Baseline** | SuperFi (Thermofisher) | 40+12 [a] | Gel | Beads | *E.coli* (DH5α) | Nextera XT | Integrase (pLAI) |
| **Q5** | Q5 (NEB) | . | . | . | . | . | . |
| **PCR free** | . | 12 | . | . | . | . | Integrase (extended) |
| **Beads** | . | . | Beads | . | . | . | . |
| **Exosap** | . | . | Exosap | . | . | . | . |
| **Gel purification** | . | . | . | Gel | . | . | . |
| **TG1** | . | . | . | . | *E.coli* (TG1) | . | . |
| **Alternative tagmentation** | . | . | . | . | . | PCR | . |
| **AmpR** | . | . | . | . | . | . | *AmpR* (pLAI) |
| **RpoB** | . | . | . | . | . | . | *RpoB* (DH5α) |
| **NextSeq (Illumina)** | . | . | . | . | . | . | . |

[a] PCR amplification was performed only as necessary by the tagmentation library preparation (methods)

**Table S2. Sequencing statistics for the homogeneous samples used to develop AccuNGS.**

| Sample | Target length | #positions called[1] | #Raw reads | #Bases called[1] | Median coverage per position[1] | Gamma 95%[2] | Median transition frequency[2] |
|---|---|---|---|---|---|---|---|
| Baseline | 1,253 | 1,232 | 2,879,486 | 237,984,201 | 174,062 | 1.08e-4 | 5.5e-5 |
| Baseline (biological replicate) | 1,253 | 1,230 | 3,020,054 | 238,477,800 | 181,858 | 1.18e-4 | 5.9e-5 |
| Alt. PCR enzyme (Q5) | 1,253 | 1,213 | 4,633,464 | 380,905,560 | 301,370 | 1.02e-4 | 5.4e-5 |
| PCR free | 1,518 | 1,502 | 4,686,406 | 368,974,699 | 243,245 | 1.03e-4 | 5.2e-5 |
| TG1 | 1,253 | 1,230 | 3,614,432 | 278,868,449 | 219,911 | 1.21e-4 | 6.3e-5 |
| Alt. PCR purification (beads) | 1,253 | 1,237 | 14,478,306 | 1,037,458,862 | 778,944 | 8.24e-5 | 4.9e-5 |
| Alt. PCR purification (Exosap) | 1,253 | 1,230 | 3,497,002 | 270,746,073 | 212,463 | 9.83e-5 | 5.05e-5 |
| Alt. tagmentation (PCR) | 250 | 250 | 337,378 | 37,381,440 | 153,097 | 1.36e-4 | 5.9e-5 |
| Alt. library purification (Gel) | 1,253 | 1,230 | 2,410,520 | 187,964,380 | 142,055 | 2.24e-4 | 5.66e-5 |
| RpoB | 1,031 | 1,011 | 2,759,920 | 203,445,548 | 199,791 | 1.11e-4 | 5.4e-5 |
| In Vitro RNA | 1,437 | 1,422 | 6,485,480 | 482,121,530 | 352,788 | 1.27e-4 | 6.8e-5 |
| AmpR (Q30) | 648 | 648 | 24,574,758 | 1,617,793,256 | 2,600,115 | 9.59e-5 | 5.2e-5 |
| AmpR (Q38) | 648 | 633 | 24,574,758 | 816,184,572 | 1,322,823 | 8.72e-5 | 4.1e-5 |
| AmpR (Nextseq, Q30) | 648 | 648 | 6,101,462 | 383,322,402 | 576,447 | 1.6e-4 | 8.1e-5 |
| pLAI gag control (Q38) | 1,860 | 1,701 | 12,824,998 | 710,846,058 | 388,422 | 8.17e-5 | 3.9e-5 |

[1] Positions were considered for this column if their coverage exceeded 10,000x.

[2] Positions were considered for this column if their coverage exceeded 100,000x.

**Table S3. Relative contribution of identified factors to AccuNGS transition errors.** Data were calculated based on the relative improvement in median error rate in each DNA control sample, compared to the baseline protocol. Relative improvement for each control was calculated by comparing the median error frequency for each transition type. Numbers in brackets refer to the p-value obtained from comparing the mutation frequencies in the altered sample against the baseline sample using two-sided Mann-Whitney U-test.

| Transition | PCR effect | Gel effect | Q-score effect | Unexplained |
|---|---|---|---|---|
| **A>G** | 2% (0.34) | 15% ($7.9 \times 10^{-10}$) | 26% ($1.4 \times 10^{-11}$) | 57% |
| **C>T** | 8% (0.016) | 8% (0.04) | 17% (0.0005) | 66% |
| **G>A** | 16% (0.001) | 11% (0.02) | 22% (0.0013) | 50% |
| **T>C** | 0% (0.33) | 14% ($6.1 \times 10^{-12}$) | 18% ($5 \times 10^{-14}$) | 67% |

**Table S4. Fitted gamma distributions for different Q-scores cutoffs and substitution types, based on the *AmpR* amplicon.**

| Q-score Cutoff | Substitution type | Number of Sites (N) | Shape (κ) | Scale (θ) | 95 percentile of fitted gamma | Mean error rate | Median error rate |
|---|---|---|---|---|---|---|---|
| **Q30** | A->G | 145 | 9.57 | 6.05E-06 | **9.17E-05** | **5.79E-05** | 5.7E-05 |
| | C->T | 173 | 3.65 | 1.40E-05 | **1.01E-04** | **5.10E-05** | 4.3E-05 |
| | G->A | 155 | 4.63 | 1.01E-05 | **8.77E-05** | **4.70E-05** | 4.5E-05 |
| | T->C | 171 | 11.08 | 5.29E-06 | **9.02E-05** | **5.86E-05** | 5.5E-05 |
| **Q38** | A->G | 142 | 5.63 | 8.31E-06 | **8.33E-05** | **4.67E-05** | 4.2E-05 |
| | C->T | 170 | 2.70 | 1.66E-05 | **9.70E-05** | **4.48E-05** | 3.55E-05 |
| | G->A | 151 | 3.28 | 1.25E-05 | **8.42E-05** | **4.12E-05** | 3.5E-05 |
| | T->C | 168 | 7.41 | 6.30E-06 | **7.80E-05** | **4.66E-05** | 4.5E-05 |

**Table S5. SuperScript III median error rates estimations.**

| From\To | A | C | G | T | Total |
|---|---|---|---|---|---|
| **A** | | 1.03E-05 | 2.69E-05 | 8.88E-06 | **4.6E-05** |
| **C** | 2.96E-05 | | 9.92E-07 | 1.17E-05 | **4.23E-05** |
| **G** | 3.43E-06 | 2.71E-06 | | 1.12E-05 | **1.74E-05** |
| **U** | 4.32E-06 | 1.09E-05 | 7.79E-07 | | **1.6E-05** |

Estimations are based on the difference in the medians of each mutation category between the In Vitro RNA control library and the AccuNGS baseline DNA protocol (pLAI). The RNA control library errors includes both the T7 RNA polymerase errors and SuperScript III errors, and therefore the provided estimates are expected to be upper bounds.

**Table S6. Summary of all clinical samples sequenced**.

| Virus | Virus sample ID | Viral load (copies/ml) | Sampling compartment |
|---|---|---|---|
| HIV | 1 | 13,000,000 | Plasma |
| HIV | 2 | 1,700,000 | Plasma |
| HIV | 3 | 700,000 | Plasma |
| HIV | 4 | 750,000 | Plasma |
| HIV | 5 | 2,400,000 | Plasma |
| HIV | 6 | 510,000 | Plasma |
| HIV | 7 | 2,280,000 | Plasma |
| HIV | 8 | 534,000 | Plasma |
| HIV | 9 | >10,000,000 | Plasma |
| RSV | 1 | 655,465 | Nose swabs |
| RSV | 2 | 430,279 | Nose swabs |
| RSV | 3 | 2,261,608 | Nose swabs |
| RSV | 4 | 3,250,291 | Nose swabs |
| RSV | 5 | 514,041 | Nose swabs |
| RSV | 6 | 256,825 | Nose swabs |
| RSV | 7 | 273,892 | Nose swabs |
| RSV | 8 | 720,462 | Nose swabs |
| RSV | 9 | 108,208 | Nose swabs |

| | | | |
|---|---|---|---|
| RSV | 10 | 94,244 | Nose swabs |
| RSV | 11 | 1,299,363 | Nose swabs |
| RSV | 12 | 1,730,379 | Nose swabs |
| RSV | 13 | 621,242 | Nose swabs |
| RSV | 14 | 108,208 | Nose swabs |
| RSV | 15 | 1,794,931 | Nose swabs |
| RSV | 16 | 550,137 | Throat swabs |
| RSV | 17 | 94,244 | Throat swabs |
| RSV | 18 | 3,183,439 | Nose swabs |
| RSV | 19 | 110,651 | Nose swabs |
| RSV | 20 | 337,658 | Nose swabs |
| RSV | 21 | 378,862 | Nose swabs |
| RSV | 22 | 59,733 | Nose swabs |
| RSV | 23 | 171,142 | Nose swabs |
| RSV | 24 | 240,283 | Throat swabs |
| RSV | 25 | 2,077,068 | Nose swabs |
| CMV | 2 | 51,575,950 | Amniotic fluid |
| CMV | 3 | 13,507,500 | Amniotic fluid |
| CMV | 4 | >8,000,000 | Amniotic fluid |
| CMV | 5 | >29,000,000 | Amniotic fluid |
| CMV | 8 | >5,000,000 | Urine |
| CMV | 9 | >7,000,000 | Saliva |
| CMV | 10 | 5,763,100 | Urine |
| CMV | 11 | >250,000,000 | Saliva |
| CMV | 12 | 2,938,000 | Urine |
| CMV | 13 | >800,000,000 | Saliva |

**Table S7. Mean expected theoretical error rates of AccuNGS.**

| Step | Error rate | Number of rounds | Expected error |
|---|---|---|---|
| Polymerase Chain Reaction (PCR) | $6.47x10^{-7}$ [Platinum SuperFi DNA polymerase by ThermoFisher] | 40 + 12[a] | $1.68x10^{-5}$ |
| Sequencing | $10^{-6}$ (Q30 filtering & overlapping paired reads) | NA | $1x10^{-6}$ |
| **Total (DNA starting material)** | | | **$1.78x10^{-5}$** |
| Reverse Transcriptase (RT) | $3.1x10^{-5}$ – $6.5x10^{-5}$ [SuperScript III/IV RT by ThermoFisher] (Potter, et al. 2003; Orton, et al. 2015) | 1 | $4.8x10^{-5}$ |
| **Total (RNA starting material)** | | | **$6.58x10^{-5}$** |

[a] An additional twelve rounds of PCR refer to PCR rounds during the tagmentation protocol (Methods)
